# Causal interpretations of family GWAS in the presence of heterogeneous effects

**DOI:** 10.1101/2023.11.13.566950

**Authors:** Carl Veller, Molly Przeworski, Graham Coop

## Abstract

Family-based genome-wide association studies (GWAS) have emerged as a gold standard for assessing causal effects of alleles and polygenic scores. Notably, family studies are often claimed to provide an unbiased estimate of the average causal effect (or average treatment effect; ATE) of an allele, on the basis of an analogy between the random transmission of alleles from parents to children and a randomized controlled trial. Here, we show that this interpretation does not hold in general. Because Mendelian segregation only randomizes alleles among children of heterozygotes, the effects of alleles in the children of homozygotes are not observable. Consequently, if an allele has different average effects in the children of homozygotes and heterozygotes, as can arise in the presence of gene-by-environment interactions, gene-by-gene interactions, or differences in LD patterns, family studies provide a biased estimate of the average effect in the sample. At a single locus, family-based association studies can be thought of as providing an unbiased estimate of the average effect in the children of heterozygotes (i.e., a local average treatment effect; LATE). This interpretation does not extend to polygenic scores, however, because different sets of SNPs are heterozygous in each family. Therefore, other than under specific conditions, the within-family regression slope of a PGS cannot be assumed to provide an unbiased estimate for any subset or weighted average of families. Instead, family-based studies can be reinterpreted as enabling an unbiased estimate of the extent to which Mendelian segregation at loci in the PGS contributes to the population-level variance in the trait. Because this estimate does not include the between-family variance, however, this interpretation applies to only (roughly) half of the sample PGS variance. In practice, the potential biases of a family-based GWAS are likely smaller than those arising from confounding in a standard, population-based GWAS, and so family studies remain important for the dissection of genetic contributions to phenotypic variation. Nonetheless, the causal interpretation of family-based GWAS estimates is less straightforward than has been widely appreciated.

## 1 Introduction

The standard genome-wide association study (GWAS) relies on a population sample to estimate the strength of associations between trait variation and loci across the genome. The approach does not only infer the direct genetic effects that are often of primary interest, however (i.e., the effects of alleles carried by a person on that person’s trait value). Instead, estimates from population-based GWASs may also include indirect genetic effects of parents and other relatives, as well as absorb genetic and environmental confounding (Vilhjálmsson and Nordborg 2013; Young et al. 2019). In the presence of these additional effects, population-based GWASs provide biased estimates of the direct genetic effects. The extent to which this bias is a concern depends on the particular application, but confounding is a clear impediment for studies aimed at identifying causal genetic mechanisms.

Because for many traits, there are a large number of GWAS associations or loci, each of which explains only a tiny proportion of variance, researchers often focus on aggregate properties of the GWAS loci, such as genetic correlations, or predict individual trait values by combining estimated effect sizes across loci into a ‘polygenic score’ (PGS). The issues of confounding can be more pronounced for these aggregate measures, as systematic biases are compounded (Berg et al. 2019; Sohail et al. 2019; Mostafavi et al. 2020; Border et al. 2022a,b).

The possible contribution of non-direct genetic effects—in particular of environmental confounding— has motivated a turn towards family-based GWASs, which overcome the limitations of population-based GWASs by taking advantage of the randomness of Mendelian segregation from parent to child (Young et al. 2019). By holding constant differences in environments among families and randomizing alleles across the genetic backgrounds on other chromosomes, family-based GWASs provide estimates that are largely robust to the contribution of non-direct genetic effects (Spielman et al. 1993; Allison 1997; Abecasis et al. 2000; Howe et al. 2022; Young et al. 2022; Veller and Coop 2023). In their reliance on randomization, family-based studies resemble natural experiments or randomized controlled trials (RCTs) (Davey Smith and Ebrahim 2003; Madole and Harden 2023a), gold standards of causal inference in the medical and social sciences.

In causal inference, the effect of interest is often that of a treatment, which can be defined in terms of a hypothetical manipulation, in which each person is moved from untreated to treated (Neyman 1990; Rubin 1974; Holland 1986; Rubin 2005). In reality, it is not possible to observe both the treated and untreated outcomes for the same person. Instead, what an RCT provides is a comparison of treated and untreated individuals who, because of the randomization procedure, are assumed to be similar in all other regards (in expectation, or asymptotically). The mean difference in outcome between treated and untreated individuals provides an estimate of the causal treatment effect, which has an interpretation in terms of the counterfactual thought experiment in which, other than the change in the treatment status, all else is held equal.

In many contexts, it cannot be assumed that the treatment effect is identical across individuals. In the presence of heterogeneous treatment effects, the question becomes whether the effect has internal validity, i.e., whether it provides an unbiased estimate of the mean effect for the entire sample—what in social sciences is known as an average treatment effect or ATE—or an unbiased estimate for a well-defined subsample, that is a “local” ATE or LATE (Imbens and Angrist 1994; Imbens 2010). For instance, in an RCT, there may be “compliers” with the treatment and “non-compliers”, and the estimate of the treatment effect is then a LATE for the compliers. A related question is the extent to which the estimate obtained from one sample generalizes to other samples from the population or to other populations, i.e., whether the estimate has external validity. Borrowing from this language, family-based GWASs are often presented as providing an ATE for causal (direct) genetic effects, either by envisaging the hypothetical swap of one or many causal alleles in the child or by invoking an analogy to RCTs, in which the treatment is the allele (or PGS value) inherited by the child and the outcome is the child’s trait value (Davey Smith and Ebrahim 2003; Morris et al. 2020; Becker et al. 2021; Madole and Harden 2023a,b; Meyer et al. 2023).

Despite the presentation of family-based GWASs as providing estimates of *average* treatment effects, there has been little or no discussion of the impact of heterogeneity in effect sizes on family-based GWAS estimates. Yet in genetics and quantitative genetics, the importance of heterogeneous genetic effects has long been recognized and, in contrast to many other contexts in which ATEs may be obtained, its possible sources are understood (notably, as arising from gene-by-environment interactions and epistasis). In plants and animals in which environments and/or genotypes can be controlled, such gene-by-environment (G*×*E) and gene-by-gene (G*×*G) effects are ubiquitous. Similarly, studies in a wide range of species have established norms of reaction, in which the phenotypic effect of a genotype depends on the context (Lynch and Walsh 1998, Ch. 22). In human genetics, where similar manipulations are obviously infeasible, the tools to study these phenomena are much more indirect, relying entirely on statistical models. Using these approaches, the evidence for G*×*E and G*×*G is limited, but there are well-known examples for specific loci, and a number of lines of evidence for varying genetic effects across environmental settings (e.g. Barcellos et al. 2018; Young et al. 2018b; Zhu et al. 2023; Hou et al. 2023; Durvasula and Price 2023). Even in the absence of G*×*E or G*×*G, heterogeneous genetic effects can arise from varying patterns of linkage disequilibrium (LD) across families. These observations suggest that genetic effects plausibly vary with the familial environment, raising the question of whether family-based GWAS estimates have internal and external validity when they do.

This question is all the more relevant because of the ways in which family data have been used to date. For most traits, sample sizes remain too small for family-based GWASs to be feasible. Instead, a PGS is built using effect-size estimates from a population-based GWAS. Then, in a second step, researchers use the data from families to test whether, and to what extent, the population-based PGS reflects direct genetic effects. Specifically, they test a null model that direct genetic effects do not contribute at all to the population PGS, by asking if the PGS is predictive of trait differences within families—that is, whether a regression of the trait value on the PGS has a non-zero slope when controlling for parental PGSs. In addition, they often quantify the extent to which the direct genetic effects contribute to variation in the PGS by assuming that the slope of this regression provides an unbiased estimate of the direct causal effect of the PGS. The validity of these procedures relies on both internal validity of the estimates produced by the family-based analyses as well as their external validity for the sample in which the standard population-based GWAS was conducted.

Here, we interrogate these assumptions. To that end, we first define the causal effect on a trait of an allele at a single locus, in terms of counterfactual manipulations at the locus. Next, we examine under what conditions a population-based or family-based association study at the locus provides an unbiased estimate of the causal effect thus defined, either in the whole sample or a well defined subset—i.e., we delimit when the estimates obtained can be considered ATEs or LATEs. We then consider aggregate effects of many loci, as combined in a polygenic score.

## 2 Results

We assume that, for each individual in a population, the genetic contribution to some trait of interest is additive across causal loci, but that there is heterogeneity in the effects of alleles across individuals. We consider that an investigator is interested in the average causal effect on the trait of some allele, or of a polygenic score, in a sample of individuals taken from the population. The investigator has access to the genotypes of the individuals in the sample as well as to the genotypes of their parents. (Alternatively, for family-based studies, they might have access to the genotypes of full siblings.)

Concerned about indirect effects as well as environmental and genetic confounding, the investigator aims to use the randomness of Mendelian segregation in families in order to more cleanly estimate the average causal effect of an allele or polygenic score on the trait in the offspring generation. They may be also interested in using the causal effect estimated in the sample to learn about the population from which the sample is drawn. They may also want to use the estimated causal effect in order to construct a genetic intrument, with the goal of identifying other causal relationships with the phenotype, as in Mendelian Randomization designs (Brumpton et al. 2020). To make things comparable, in discussing family GWASs, standard GWASs, or hypothetical manipulations, we assume that we are measuring phenotypic outcomes in the children’s generation.

### 2.1 The effects of alleles at a single locus

To generate intuition for the influence of G***×***E (and G***×***G) interactions on population and family-based GWAS estimates, we initially focus on a single bi-allelic locus, at which the two alleles have a direct causal effect on the trait of interest. We later consider the case where the alleles at the genotyped locus do not causally affect the trait, but instead tag alleles at a nearby causal locus. Label the two alleles *A*_1_ and *A*_2_, with *A*_2_ the “focal” allele. We assume that there is no genetic or environmental confounding, and no indirect genetic effect via relatives or peers, making possible what we henceforth call an “unconfounded GWAS”. To incorporate G*×*E interactions, we allow the effect sizes of the alleles at the locus to depend on the family environment. The phenotype of individual *i* in family *f* is

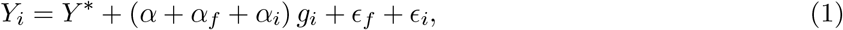

where *g*_*i*_ is the number of copies of the focal allele *A*_2_ carried by individual *i* at the locus (0, 1, or 2), and *ϵ*_*f*_ and *ϵ*_*i*_ are family- and individual-specific environmental deviations in the trait’s value, with mean zero. *Y* ^*\**^ is an intercept value of the trait. *α* is the mean genetic effect of the focal allele. The family- and individual-specific deviations of the genetic effect due to G*×*E are *α*_*f*_ and *α*_*i*_; we define their population means to be zero: 𝔼 [*α*_*f*_] = 𝔼[*α*_*i*_] = 0.

Given this phenotypic model, we consider two questions. First, what is a sensible *definition* of the causal effect of allele *A*_2_ on the phenotype? Second, how do the estimates obtained by a population-based or family-based association study compare to this definition of a causal effect?

#### Causal effects via manipulation

It is common to define causal effects in terms of counterfactual manipulations (Rubin 1974). To this end, we consider two thought experiments, one a randomization and one a more precise genetic manipulation.

We begin with a simple thought experiment in which we randomly reassign genotypes at the locus to individuals in the sample, independent of their environments or genetic backgrounds. This thought experiment resembles an RCT. Since, in expectation, there is zero covariance between *g*_*i*_ and *α*_*f*_ or *α*_*i*_, the expected effect of *A*_2_ is simply *α*. We note that some existing definitions of the causal effect of an allele in the presence of G*×*E in fact amount to this thought experiment (Lee and Chow 2013, Eq. 15; see Appendix A1.3)

In turn, to define a causal effect of *A*_2_ via a counterfactual genetic manipulation, we choose, randomly among the gametes produced by the parental generation, one that carries the *A*_1_ allele, and we imagine flipping the allele in this gamete to *A*_2_ (as if by CRISPR editing). If, among parents, *p* is the frequency of the focal allele *A*_2_ and *p*_11_, *p*_12_, and *p*_22_ are the three genotype frequencies, then with probability *p*_11_*/*(1*−p*) the chosen *A*_1_-bearing gamete derives from an *A*_1_*A*_1_ parent, while with probability *p*_12_*/*2(1 *−p*), it derives from an *A*_1_*A*_2_ parent. The expected difference between the resulting offspring’s phenotype if we were to flip the allele versus if we do not is therefore

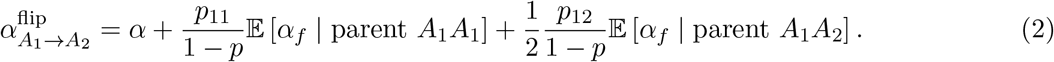

We can similarly define the causal effect of *A*_1_ as the expected phenotypic difference caused by randomly selecting an *A*_2_-bearing gamete and flipping the allele to *A*_1_:

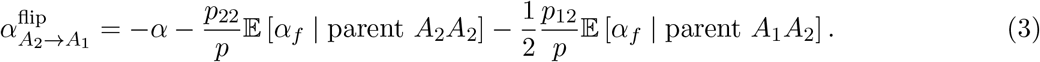

In general, the definitions (2) and (3) need not be equal in magnitude, nor need they equal *α*. The reason is that each hypothetical allele flip occurs in the original environment of the genotype. For example, if we flip *A*_1_ *→ A*_2_ in a gamete, the resulting offspring gains an *A*_2_ allele but still grows up in the original environment in which the *A*_1_ allele was originally found. If *A*_1_ and *A*_2_ alleles have systematically different distributions of interacting environments, the two manipulations (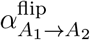 and 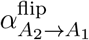) will differ in their effects and differ from *α*.

We can define a counterfactual manipulation that effectively randomizes alleles across environmental backgrounds and thus reconciles the randomization and allele-flipping definitions of the causal effect. We first note that the expected effect of the *A*_1_*→A*_2_ flip defined above, 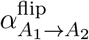, is the same, though opposite in sign, as that of flipping *A*_1_*→A*_2_ in the environments experienced by *A*_1_ alleles. Therefore, if we calculate the expected difference in an offspring’s phenotype caused by choosing a gamete at random and flipping its allele, polarizing the difference by the allele that we flip, we obtain

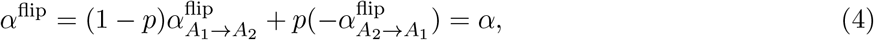

(see Appendix A1.1). Thus, we can define a sensible weighted average of the two allele-flipping effects that returns *α* as the average causal effect at the locus. Note that *α* is an average effect for the sample, and so is a property of the specific environments experienced by the sample and their genetic backgrounds.

#### Effect-size estimates from association study designs

Next we consider whether various GWAS designs provide an unbiased estimate of the average causal effect. The genotype of each offspring can be written as the average of their parental genotypes (*g*_*m*_ + *g*_*f*_)*/*2 plus a zero-mean term *ς* that accounts for the randomness of segregation in transmissions from the parents:

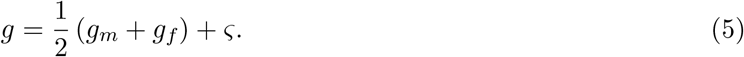

If we perform a family-based association study by regressing the trait values of offspring on their genotypes at the locus, controlling for their parental genotypes, we obtain an effect-size estimate for allele *A*_2_ of

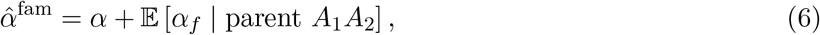

where the second term—the deviation of the family-based estimate 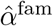 from *α*—is the average family deviation conditional on a parent being heterozygous at the focal locus (Appendix A1.2). This result is intuitive: parent-offspring studies rely on contrasting the associations of transmitted and untransmitted alleles with the phenotype of the offspring (Kong et al. 2018). When parents are homozygous, no such comparison can be made, since their transmitted and untransmitted alleles are the same. The family-based estimate therefore makes use only of heterozygous parents. In other words, we can define the family-based estimate at a single locus as a LATE in the subpopulation of children who are the offspring of heterozygous parents; in the language of a causal inference theory, those are our “compliers”. If heterozygous parents are non-randomly distributed across environments, however, the estimate obtained from a family-based GWAS may not be an unbiased estimate for the whole sample, which includes children of homozygous parents, and may not have external validity for the sample used in the standard population-based GWAS.

We note that the estimate in Eq. (6) is the same, in expectation, as would be obtained in a sibling study where the phenotypic differences between full siblings are regressed on their genotypic differences at the locus, since both designs amount to regressing offspring phenotypes on their segregation deviations *ς* (again, assuming no indirect effects of siblings; Appendix A1.2; also see Trejo and Domingue 2018; Fletcher et al. 2021).

If we instead perform a population-based association study by regressing the trait values of the offspring on their genotypes, without controlling for the parental genotypes, then, in an unconfounded GWAS, we obtain an effect-size estimate that can be written in the form

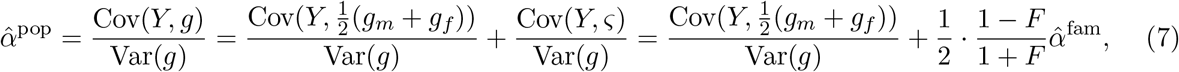

where *F* is the inbreeding coefficient at the locus, which, along with the frequency *p* of *A*_2_, we have assumed to be the same among offspring and their parents (Appendix A1.2). The first term on the right-hand side of Eq. (7) is the contribution of the genotypic variance *among* families to the population slope, while the second term is the contribution from the genotypic variance *within* families (due to random segregation). The first term (and, more generally, Eq. 7 written out explicitly) is a complicated weighted sum of the effects of *A*_2_ in the environments experienced by the offspring of the three possible parental genotypes (Appendix A1.2).

In general, 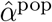 differs from the causal effect of the alleles at the locus, as defined in our hypothetical manipulation above (Eq. 4), because the association study examines the effects of the alleles at the locus in the particular environments in which they are found.

#### Summary

In the presence of heterogeneous allelic effects at a locus, the effect-size estimates produced by family- and population-based association studies need not be the same on average, nor will they equal, in expectation, the causal effects of the alleles defined via our hypothetical experimental manipulations. The underlying reason is that the quantities produced by these study designs and definitions are averages of allelic effects taken across different distributions of environments.

In Figure 1, we illustrate with a simple example how G*×*E can lead population- and family-based association studies to produce different estimates when the study sample is drawn from two populations inhabiting different environments.

**Figure 1:**
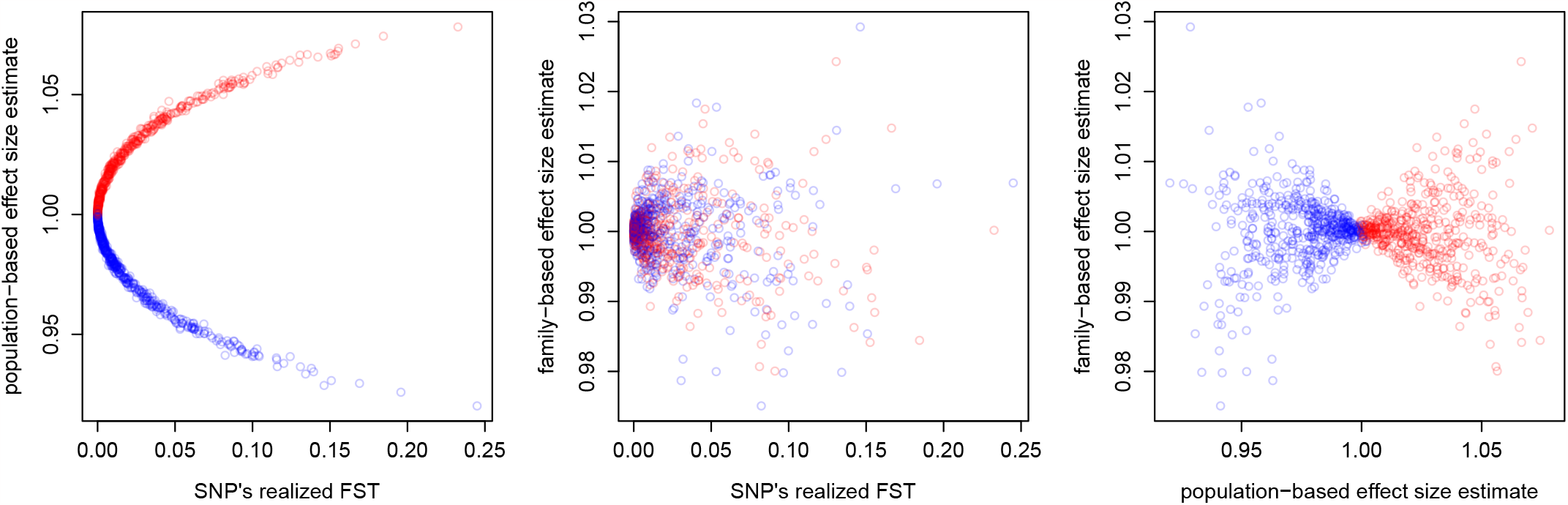
Relationships between the effect-size estimates produced by population- and family-based association studies and *F*_*ST*_ in a simple model of population structure. There are two populations, red and blue, each inhabiting a distinct environment. At a bi-allelic locus, the focal allele has effect 1.1 on a trait of interest in the environment of the red population and effect 0.9 in the environment of the blue population. We simulated independent drift of allele frequencies at the locus in the two populations, and then conducted population- and family-based association studies of the trait in a large sample drawing equally from the two populations. **A**. When drift results in some differentiation between the populations (*F*_*ST*_ *>* 0), the effect-size estimate from a population-based GWAS is systematically biased away from the true population-average effect of 1, with the direction of the bias reflecting whether the focal allele ended up at higher frequency in the red population (red points) or the blue population (blue points). **B**. The estimate from a family-based GWAS depends instead on whether heterozygosity ends up being higher in the red population or the blue population; that is, on the population in which the focal allele at the locus moved closer to frequency 1*/*2. **C**. The family-based estimate differs from the population-based estimate in the presence of G*×*E. See Appendix A1.4 for details.

#### LD differences

In practice, a locus with a signal of association in a GWAS will often not affect the trait itself. Instead, it will be in linkage disequilibrium with—and thus ‘tag’—one or more nearby loci that causally affect the trait. If patterns of LD between the marker and causal loci differ across individuals, then, even in the absence of G*×*E at the causal loci, the effects estimated for the marker locus can differ between study designs in ways that resemble those of G*×*E at the marker locus.

Consider a model with two loci, a genotyped marker locus that does not causally affect the trait of interest and an ungenotyped causal locus that does. The alleles at the marker locus are *m* and *M*, and the alleles at the causal locus are *a* and *A*. Here, we assume that there is no G*×*E at the causal locus: the effect of *A* is to increase the trait’s value by *α* in expectation, independent of the environmental setting.

First, we consider a definition of the counterfactual effect of allele *M* at the marker locus analogous to the counterfactual definition of the causal effect of the allele *A* at the causal locus, laid out above. To this end, we imagine that the investigator can swap haplotypes of a given physical length around the genotyped marker: specifically, that they can randomly choose a gamete that carries the *m* allele and flip its haplotype to a random haplotype in the sample containing *M* . If the length of the flipped haplotype is too short to contain the causal locus, the effect of this flip is simply 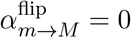, since the alleles at the marker locus do not affect the trait. However, if the flipped haplotype is long enough to include the causal locus, then the swap might change the allele at that locus too—either *a→A* or *A→a*—and therefore affect the phenotype. The manipulation effect will then depend on the proportion of marker allele *y*-containing haplotypes that also contain causal allele *x, p*_*x*|*y*_ ; the average effect of the flip is

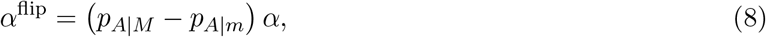

as shown in Appendix A1.5. Since we are assuming that there is no G*×*E at the causal locus, the average phenotypic effect of the reverse *M→m* haplotype switch would be the same (though opposite in sign) as Eq. (8), which is why we have omitted the *m→M* subscript in Eq. (8). While Eq. (8) is defined in terms of a single causal locus, it naturally generalizes to the case where the marker locus tags multiple causal loci within the flipped region.

In the absence of confounding, a population-based association study at the marker locus returns an effect-size estimate that is equal to the allele-flipping effect in expectation:

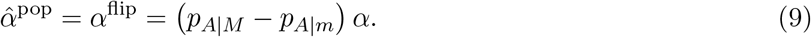

(Note that the quantities in Eqs. (8) and (9) can also be written in terms of coefficients of linkage disequilibrium—see Appendix A1.5.)

In contrast, a family-based association study at the marker locus returns an effect-size estimate of

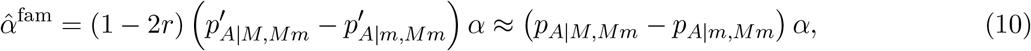

where *r* is the recombination fraction between the marker and causal loci, *p*′_*A*/ *M,Mm*_ and *p*_*A*|*M,Mm*_ are the proportions of *M* containing haplotypes that also contain *A* in heterozygous *Mm* parents and offspring, respectively, and *p*′_*A*/ *M,Mm*_ and *p*_*A*|*m,Mm*_ are analogously defined for allele *m* (Appendix A1.5). The approximation in (10) holds under the assumptions that the marker and causal loci are tightly linked (*r ≈* 0) and that genotype frequencies are similar in the parental and offspring generations.

In summary, when the marker and causal loci are tightly linked, the family-based study returns an unbiased estimate of a causal effect defined in an analogous way to Eq. (8), namely flipping haplotypes only in gametes produced by parents heterozygous at the marker locus. In this context too, therefore, the family-based estimate can be interpreted as a LATE for those offspring.

We can rewrite the family-based estimate as

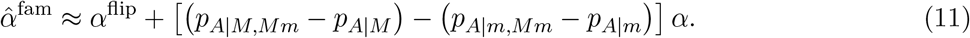

Therefore, the family-based estimate at the marker locus will differ from the causal effect defined by the allele flip (and from the effect estimated in an unconfounded GWAS) if the *A* allele tends to co-occur on the same haplotype as the *M* allele more frequently in *Mm* heterozygotes than in the rest of the population.

This phenomenon is mathematically analogous to the case of G*×*E considered above—compare Eqs. (11) and (6)—even though the genetic mechanisms are distinct.

As a simple illustration of how LD differences between heterozygous and homozygous parents can influence the various association study designs, consider two populations that differ in both the degree of LD between and the allele freqencies at the marker and causal loci. In a sample drawn from across these two populations, the effect size estimated at the marker locus by a family-based association study will more strongly reflect the degree of LD in the population with greater heterozygosity at the marker locus, whereas an unconfounded population-based association study will instead reflect differences in the haplotype frequencies across the two populations (Appendix A1.6).

### 2.2 Polygenic scores

To study the influence of G*×*E in applications of polygenic scores (PGSs), we focus on the general case of the relationship between a PGS for trait X and the value of trait Y. As a special case, X and Y could be the same trait, or the same trait but in a different set of environments.

An investigator has access to effect-size estimates 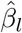 at a set of genotyped loci *l ∈* Λ from a prior GWAS on trait X. The sample across which this GWAS was carried out does not overlap with the sample used in the investigator’s subsequent analysis. Given individual *i*’s genotype *g*_*il*_, *l ∈* Λ, the investigator calculates a trait-X PGS for the individual:

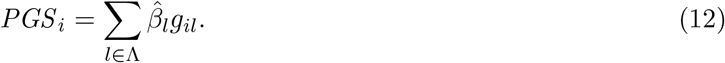

If individual *i* is in family *f*, their value for trait Y is given by

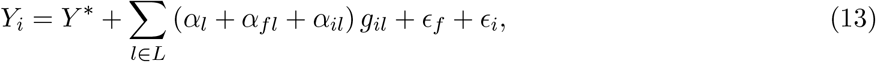

which is the multi-locus version of Eq. (1), with the set of loci *L* causally underlying variation in trait Y. We are interested in the causal relationship between the PGS for the trait X and the value of trait Y, both in terms of a hypothetical manipulation and in terms of estimates produced by standard least-squares regression approaches. We consider population-based designs that regress *Y* on *PGS* and family-based designs that regress *Y* on *PGS* controlling for the maternal and paternal PGSs.

In practice, a PGS is constructed using estimated rather than true effect sizes. Specifically, it reflects the signal of causal effects partially (and noisily) captured by the genotyped loci on which the PGS is based. Additionally, the effect-size estimates may be biased due to confounding in the GWAS. We ignore these complications here. For simplicity, we further assume that the set of loci included in the polygenic score is a subset of the loci that causally affect the trait of interest (Λ *⊆ L*) and that the causal loci are in Hardy-Weinberg and linkage equilibrium. As we show, even under these simplifying assumptions, interpreting the effects of polygenic scores in the presence of G*×*E is not straightforward. Our results extend naturally to the case where the PGS loci are not themselves causal but instead tag nearby causal loci (a point that we return to below).

#### Causal effects via manipulation

As in the single locus case, we can again consider two thought experiments to understand casual effects in the PGS case, the first involving randomization and the second involving genetic manipulation. First, we imagine randomly assigning genotypes at the loci in Λ to our sampled individuals, independent of their environments or genetic backgrounds. Under this randomization, least-squares regression of trait Y on the trait-X PGS returns, in expectation, a coefficient

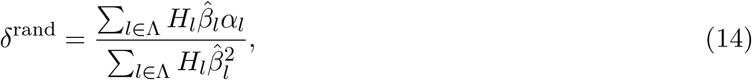

where *H*_*l*_ = 2*p*_*l*_(1 *− p*_*l*_) is the heterozygosity at locus *l* under the assumption of Hardy-Weinberg equilibrium. Under our assumption that the loci in Λ are causal for trait X, the numerator in Eq. (14) can be interpreted as the genic covariance between the trait-X PGS and trait Y, while the denominator is the genic variance of trait X.

It is more complicated to define the causal effect of the PGS in terms of hypothetical genetic manipulations, in which we imagine experimentally flipping alleles at the PGS loci: with many loci contributing to the PGS, a desired change in the value of the trait-X PGS can be achieved via many different manipulations, and these different manipulations will, in general, not have the same average effect on trait Y. However, in Appendix A2, we describe two sensible manipulation strategies that respect the sample variance structure of the PGS and return an effect of the PGS, *δ*^flip^, that is the same as that given by the randomization thought experiment described above, *δ*^rand^. Thus, Eq. (14) offers a natural way to define the causal effect of the PGS.

When the PGS loci do not themselves causally affect the trait, but instead tag ungenotyped causal loci, we can still define a causal effect of the PGS via manipulation, just as we did for the case of a single marker locus above, with the investigator flipping *haplotypes* of a given length around each PGS locus.

Under this definition, the *α*_*l*_ terms in Eq. (14) would be replaced by expressions for the average effects of the haplotype swaps (i.e., Eq. 8, assuming that the flipped haplotype is long enough at each marker locus to include the causal locus).

#### Estimates of the effect of the PGS

Family-based designs for estimating the effect of the PGS rely on the regression

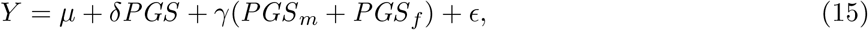

where *PGS* is the PGS of the offspring and *PGS* _*m*_ and *PGS* _*f*_ are the PGSs of the mother and father, respectively. The estimate 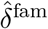 produced by OLS estimation of this regression has been interpreted as an estimate of the direct effect of the PGS on the phenotype, under the assumption that the parental PGSs effectively control for population structure and for indirect effects of genotypes of relatives on the phenotype of the offspring.

Since we can write

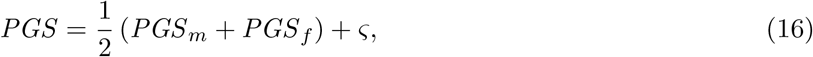

with *ς* the segregation deviation of the offspring’s PGS from the midparent value, the estimate 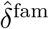 produced by OLS estimation of Eq. (15) is the same, in expectation, as that produced by OLS estimation of the regression

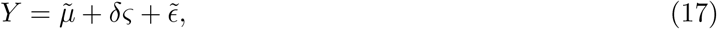

by the Frisch–Waugh–Lovell theorem (Angrist and Pischke 2009, pp. 35-36; Appendix A3.1).

In the presence of G*×*E interactions, and under the simplifying assumptions laid out above concerning the sets of PGS and causal loci Λ and *L*, the family-based estimate is, in expectation,

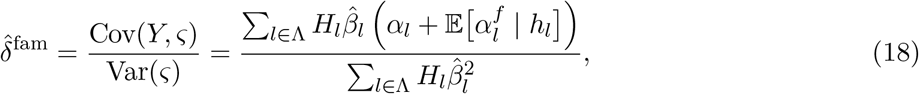

where *h*_*l*_ denotes that a parent is heterozygous at locus *l* (Appendix A3.2).

As this expression makes clear, the estimate from a family-based GWAS will differ systematically from the randomization or allele-flipping effect of the PGS if parents heterozygous at any locus in the PGS are non-randomly distributed across environments 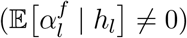.

Without information about parental genotypes, a population-based study of the effect of the PGS effect is based on the regression

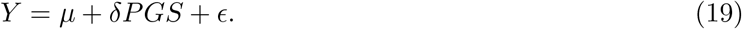

In addition to our assumption that there is no genetic confounding in the sample—that is no long distance LD between PGS loci and other causal loci—we also make a further ‘best case’ assumption that there is no environmental confounding, i.e., no covariance between environmental effects (*ϵ*) and the PGS. In that setting, from Eq. (16), OLS estimation of this regression produces

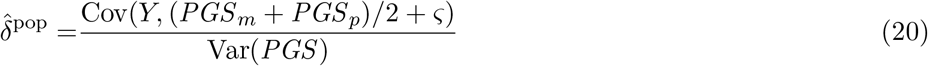

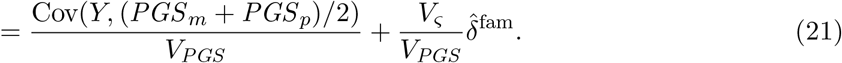

Even in the absence of genetic or environmental confounding, the estimated effect of the PGS is a mixture of between- and within-family effects. Thus, while the family-based estimate 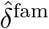 can be seen as a component of the population-level estimate 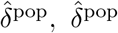, will additionally reflect the G*×*E effect in the children of parents homozygous at PGS loci.

#### Summary

In the presence of G*×*E, the family-based estimate of the causal effect of the PGS, 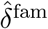, can be systematically biased away from the causal effect defined by genotype-environment randomization or by experimental manipulation of genotypes if, for any of the loci in the PGS, heterozygotes and homozygotes are distributed differently across environments. In such cases, the family-based estimate 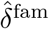 is a strangely weighted average across loci and across families (Eq. 18).

At any given locus, there exists a set of individuals, i.e., the offspring of heterozygotes, for whom a family-based GWAS provides an unbiased estimate of an average causal effect, i.e., a LATE (cf. Eq. 6). However, this set of individuals will be different for each SNP, and so, in general, 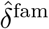 cannot be interpreted as an unbiased estimate of an average effect of the PGS for any subset of sampled individuals. In general, in the presence of an interaction between genetic effects and familial environments, family-based estimates 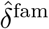 do not have internal validity.

These conclusions extend to the case where the PGS loci are not causal themselves, but instead tag nearby causal loci. In this case, the within-family estimate of the PGS effect will reflect the level of LD in heterozygotes for each marker (Eq. 11, assuming no G*×*E at causal loci). If the patterns of LD differ between heterozygous and homozygous parents, the within-family estimate is not, in general, an unbiased estimate of the average effect of the PGS defined in terms of haplotype manipulation, nor can it be interpreted as a LATE for any subsample.

#### Under what conditions is the family-based regression coefficient on the PGS an internally valid estimate of an average causal effect?

The within-family regression slope of the PGS, 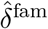, has been described as providing an estimate of the direct causal effect of the PGS. As our results show, when genetic effects interact with the familial environment or when LD patterns differ between parents that are homozygous vs. heterozygous for PGS loci, this claim does not hold. The within-family regression slope of the PGS has also been described more specifically as “a weighted average over the direct effects of the PGI for the individuals in the population”, with this average taken “over any heterogeneity of effects across individuals that may exist” (Okbay et al. 2022, SI Section 7.1). While, by the same token, this claim cannot be generally true, here we consider under what restricted conditions it is valid.

There are two special conditions that jointly allow us to recover an interpretation of the family-based regression coefficient 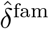 as a weighted average of direct PGS causal effects. The first is that the G*×*E effects 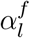 are, for each family *f*, a fixed multiple *C*_*f*_ of the population-average effect *α*_*l*_ across all causal loci, such that a family’s environment either amplifies (*C*_*f*_ *>* 0) or dampens (*C*_*f*_ *<* 0) the genetic contribution to the trait by a constant factor across loci. The second is that the effect-size estimates 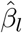 used to construct the PGS differ in expectation from the population-average causal effects *β*_*l*_ only by a multiplicative factor that is constant across loci (allowing also for uncorrelated noise).

If these two conditions both hold, each family is characterized by a well-defined PGS slope *δ*_*f*_ = (1 + *C*_*f*_)*/B*, such that any genetic manipulation that increases the PGS of an offspring in family *f* by 1 unit—no matter which loci are flipped and in what direction—increases the offspring’s trait value by *δ*_*f*_ in expectation. The regression coefficient 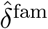 can then be interpreted as an average of these family-specific slopes *δ*_*f*_ .

Even under these strict conditions, however, this average does not weight each family equally, and therefore does not return the average PGS slope in the sample. Instead, each family is weighted proportionally to its segregation variance for the PGS, with families in which parents are heterozygous at more of the PGS loci upweighted in the calculation of 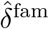 (Appendix A3.3). The reason is that these families contribute more within-family variation in the PGS, and it is within-family PGS variation upon which family-based estimation of *δ* depends. Thus, while the estimate of *δ* produced by the within-family PGS regression is not an ATE across the sample of genotyped families, it is, under the specific conditions laid out above, a LATE in an “effective sample” (Aronow and Samii 2016) of families weighted according to the parents’ PGS segregation variance. This is a specific example of a more general phenomenon, known in statistics and econometrics, where, in estimating of a coefficient of interest, controlling for additional covariates (in our case, the parental PGSs) causes OLS to upweight some observations and downweight others (e.g., Angrist and Krueger 1999; Aronow and Samii 2016; Angrist and Pischke 2009, pp. 76, 79; see Appendix A3.3).

While these parallels to regression results from other fields are enlightening, in practice, the assumptions of a fixed multiplicative G*×*E effect across loci and a fixed multiplicative bias in the effect-size estimates used to construct the PGS are unlikely to be met. When considering the PGS as a “treatment”, we rely on a sum of small estimated effects. Because the causal loci tagged by the PGS loci will often be pleiotropic in their effects (i.e., involved in multiple biological processes), the property of a single G*×*E multiplier *C*_*f*_ —although a useful heuristic—is unlikely to hold for the trait of interest in the family-based study. Moreover, it is worth noting that the estimated weights used to build the PGS are not just noisy but often systematically biased in heterogeneous ways across loci.

#### Is there a causal interpretation of the within-family PGS slope?

After computing the within-family slope of the PGS, 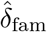, some family-based studies proceed to interpret the fraction of the sample-wide phenotypic variance *V*_P_ that the fitted values explain, 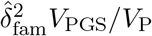, as an estimate of the proportion of phenotypic variation that is due to direct causal effects of (or tagged by) the PGS loci (Meyer et al. 2023). In light of our arguments above, this interpretation is not valid under general conditions.

The estimate 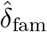 is estimated from—and therefore only strictly applies to—the within-family PGS variance *V*_*ς*_ . Attributing the explanatory value of 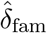 to the full population PGS variance *V*_PGS_ requires an extrapolation from within to between families. As we have shown, the within-family slope generally lacks internal validity, and so this extrapolation can lead to a biased estimate of the variance explained by direct effects in the presence of heterogeneity. Similar biases may also affect other methods that use genetic segregation from parents in order to partition the population phenotypic variation into genetic and non-genetic components (Visscher et al. 2006; Young et al. 2018a).

The population variance of the PGS can be decomposed into contributions from within and between families:

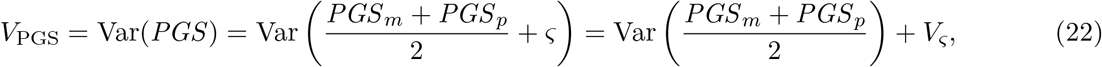

noting that the segregation deviation *ς* is uncorrelated with (*PGS* _*m*_ + *PGS* _*p*_)*/*2. If we focus on the within-family variance *V*_*ς*_, and consider the proportion of the overall phenotypic variance explained by fitted values 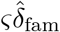 based on it, we obtain

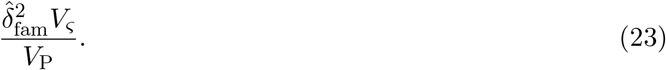

This approach is internally coherent, with the fraction properly interpretable as the contribution to phenotypic variance arising from Mendelian segregation within families at the PGS loci. This interpretation is valid because families are weighted the same in the calculation of 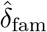 as they are in *V*_*ς*_ ; the interpretation of Eq. (23) therefore does not require an extrapolation from families with greater segregation variance (more heterozygous parents) to families with lower segregation variance (less heterozygous parents).

Eq. (23) can also be interpreted in terms of an imaginary manipulation that would produce a quantity measureable at the population level. We first note that variance due to Mendelian inheritance from parents at the PGS loci can be set to zero if each individual’s phenotype is replaced by their phenotype averaged over the random inheritance of their parental genotypes at the PGS loci, holding all the other genotypes and environments constant. One might imagine, for example, a hypothetical manipulation in which an investigator measures the mean conditional phenotype over a large number of individuals whose genotypes are identical to the individual but in which alleles (or haplotypes) at the PGS loci that are heterozygous in the parents have been flipped in the zygotes and the children raised in exactly the same environment as the original individual. The experimental replacement of individuals by their conditional means would reduce the population variance by 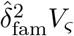, and thus proportionately 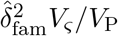.

These results establish a solid basis for causal statements about the proportion of phenotypic variance explained by within-family PGS variance. In the absence of assortative mating or other sources of long-range signed linkage disequilibrium, the within-family variance of the PGS is expected to be half the sample-wide variance (*V*_*ς*_ = *V*_PGS_*/*2) (less if alleles with the same directional effect on the trait are in positive LD, as is the case with assortative mating, and more if they are in negative LD, as arises under various kinds of selection). Therefore, our results indicate that only half (and potentially less) of the phenotypic variance attributed to the PGS based on within-family estimates of its effect can potentially be estimated in an unbiased manner, while the rest relies on an extrapolation based on the assumption of no heterogeneity of genetic effects across families.

A caveat is that we have not considered the multiple covariates often included in family studies beyond parental genotypes, which will further complicate the interpretation of causal effects (Imbens and Wooldridge 2009). Clearly, further work is required to define precisely what is being estimated, in practice, in family-based GWASs.

## 3 Discussion

While the random inheritance of parental alleles by children offers a natural experiment (Fisher 1952), the genetic variation that this experiment generates—based on which one hopes to establish “treatment” effects of alleles on phenotypes of interest—is contributed only by heterozygous parents. In other words, we do not observe alternative allelic treatments among the offspring of homozygotes and therefore cannot learn about treatment effects on their phenotypes. In contrast, the effect of an allele across families in a population reflects inheritance from both homozygous and heterozygous parents. The consequence is that if there is heterogeneity in effects between the children of homozygous and heterozygous parents, family studies will result in a biased estimate of the average effect of an allele in a population. In the case of the effect size estimated by a family GWAS for a single locus, the estimate can nonetheless be viewed as a LATE for the children of heterozygotes, and thus has internal validity for a well-defined subset of families. The same does not hold for a PGS, however, which has no general interpretation as a LATE at the population level in the presence of effect-size heterogeneity.

That so many fields of study focus on reliably estimating *average* treatment effects reflects the fact that heterogeneity in causal effects is ubiquitous. Genetics is no exception, with many well studied sources of heterogeneity, including G*×*G and G*×*E interactions, and for marker loci, varying patterns of LD. For the specific issues with family studies raised here to be a problem, however, requires two conditions to be met: (i) genetic effects must be heterogeneous across environments and (ii) genotypes must be non-randomly distributed across the relevant environments. Given extensive evidence for heterogeneity in the effects of alleles due to G*×*E (and G*×*G) in settings where experimental manipulations are possible, the first condition seems highly plausible. In turn, the evidence of environmental and genetic confounding in human GWAS studies for an increasing number of traits shows that SNP genotypes are non-randomly distributed across environmental and genetic backgrounds. Thus, it seems important to consider what happens to estimates of family-based GWASs in this setting.

In doing so, we have focused primarily on G*×*E as the source of heterogeneity in genetic effects. As noted above, however, the effect of an allele could also systematically differ across families (*α*_*f*_) because it is involved in epistatic interactions with alleles at other loci in the genome (i.e., because of G*×*G). By analogy to our G*×*E model above, epistatic interactions would lead to biases in family-based GWASs if parents who are heterozygous at the focal study locus tend to have systematically different genotypes at loci that interact epistatically with the focal locus, relative to the population distribution of such genetic backgrounds. We have also ignored parent-offspring interactions in family-based studies. Following the same logic, interactions between alleles of parents and offspring will result in family GWAS estimates that are the average effect of the focal allele in an offspring conditional on the genetic background of a heterozygous parent. Thus, a non-random distribution of genetic backgrounds in heterozygous parents is also a potential source of bias.

In addition to interactions with the environment or other loci, differences between homozygotes and heterozygotes in the degree of LD between marker loci and the causal loci they tag can also give rise to variation in effect sizes at the marker loci. As we have shown (Eq. 11), these differences can also lead to biases in family-based estimates of average effect sizes. One plausible way that such differences in LD can arise is when the GWAS draw samples from across ancestry gradients; for example, from across the north-south gradient in haplotype diversity within Europeans (Auton et al. 2009). Evidence that changes in LD patterns contribute to differences in the prediction accuracy of PGSs over even short genetic distances (Wang et al. 2020; Privé et al. 2022; Ding et al. 2023) supports the view that sampling families from across ancestry gradients may be an important source of heterogeneity.

To provide an intepretation of the effect size being estimated at a tagging—rather than causal—locus, we proposed a thought experiment in which a haplotype of a given length is experimentally swapped. Under the assumption of tight linkage between the causal and marker locus and no LD between causal loci, the effect estimated in family studies corresponds to the average effect of this manipulation in the offspring of heterozygotes at the marker locus. In the presence of LD between causal loci, the family-based effect-size estimate at the marker locus absorbs the effects of any causal loci on its chromosome that are in LD with it, complicating even the single-locus interpretation (Veller and Coop 2023). More generally, we have not considered how LD among causal loci will complicate causal intepretation of polygenic scores. As our results show, even in the simpler case of uncorrelated causal loci, articulating a causal interpretation of family-based studies is not straightforward.

Further complications will arise if there is selection bias on the participants in the population-based or family-based GWAS (Benonisdottir and Kong 2023), as the effects estimated may then be biased relative to those of a representative sample. An additional issue specific to family studies is that the key assumption of allele randomization by Mendelian segregation will be violated when the selection of families in the study occurs based on a heritable phenotype.

## Conclusion

To date, one use of family studies has been to provide rigorous evidence that the PGS contains *some* signal of a genetic causal effect. While this use is valid, one might argue that, given that it is well established that almost any trait is partially heritable and that genetic correlations are widespread, it should not be surprising that there are non-zero genetic effects on a trait (Barton and Keightley 2002; Turkheimer 2000). The focus on family studies providing ATEs reflects the desire for richer quantitative answers. Our results indicate that, in a general setting, a number of claims about what can be learned from family-based GWASs about population differences are not on a solid causal footing. However, our results also establish that family studies can be interpreted as providing an unbiased estimate of the population contribution of segregation of PGS alleles *within* families (Eq. 23). Since, under standard quantitative genetics models, the within-family variance is a sizable contribution to the population variation, even this more limited used of family-based GWASs may yield important insights. Moreover, in practice, the biases that arise from family studies due to various sources of heterogeneity are likely smaller than the effects of confounding on population-based GWAS estimates. In that sense, estimates from family studies are more interpretable as causal effects than those from population-wide studies, and discrepancies between family- and population-based GWASs still offer a useful heuristic for identifying and dissecting how confounding affects PGSs.

## Acknowledgements

We thank David Blei and Cyrus Samii for helpful discussions, and Vince Buffalo, Doc Edge, and Arbel Harpak for comments on an earlier draft of the manuscript. Funding was provided by the National Institutes of Health (NIH R35 GM136290 awarded to GC and R01 HG011432 co-awarded to MP) and a Branco Weiss fellowship to CV.

## A1 Single-locus effects

In all of our calculations, we assume that the investigator has access to the genotypes of two generations: parents and offspring.

We first consider the simple case where the alleles at a given genotyped locus causally affect a trait of interest. At the locus, there segregate two alleles, *A*_1_ and *A*_2_, with *A*_2_ designated the ‘focal’ allele. The relationship between genotype and phenotype among offspring is

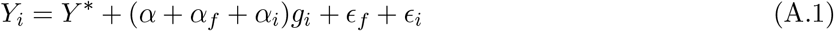

for individual *i* in family *f*, where *g*_*i*_ is the number of copies of *A*_2_ that individual *i* carries at the locus, *ε*_*f*_ and *ε*_*i*_ are zero-mean deviations that are independent of *g*_*i*_ (no environmental or genetic confounding, and no indirect effects), and 𝔼 [*α*_*f*_] = 𝔼[*α*_*i*_] = 0. *Y* ^*\**^ is the expected value of the trait for an *A*_1_*A*_1_ homozygote.

The frequency of the focal allele *A*_2_ is *p*_2_ = *p*, while the frequency of *A*_1_ is *p*_1_ = 1 *− p*. The frequencies of the three genotypes, *A*_1_*A*_1_, *A*_1_*A*_2_, and *A*_2_*A*_2_ are *p*_11_ = (1 *− p*)^2^ + *p*(1 *− p*)*F, p*_12_ = 2*p*(1 *− p*)(1 *− F*), and *p*_22_ = *p*^2^ + *p*(1 *− p*)*F*, where *F* is the inbreeding coefficient at the locus. We assume these frequencies to be constant across the parental and offspring generations. The genotypic variance at the locus is Var(*g*_*i*_) = 2*p*(1 *− p*)(1 + *F*).

### A1.1 Defining the causal effects of *A*_1_ and *A*_2_ via counterfactuals

We first calculate the causal effects of *A*_2_ and *A*_1_ defined in terms of counterfactual experimental allele swaps, as set out in the Main Text. For the effect of *A*_2_, we randomly choose a parental gamete containing *A*_1_ and flip the allele to *A*_2_. With probability *p*_11_*/p*_1_ = 1 *− p* + *pF*, the parent who produced the gamete is *A*_1_*A*_1_, and with probability *p*_12_*/*(2*p*_1_) = *p*(1 *− F*) they are *A*_1_*A*_2_. The average effect of the flip on the offspring’s phenotype is therefore

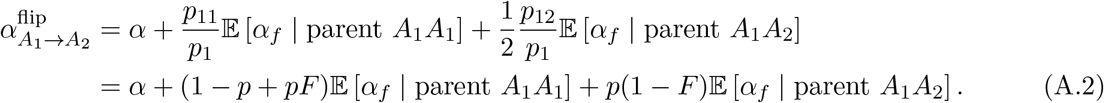

For the analogous definition of the causal effect of *A*_1_, we randomly choose a gamete containing *A*_2_ and flip it to *A*_1_. With probability *p*_22_*/p*_2_ = *p* + (1 *− p*)*F*, the parent who produced the gamete is *A*_2_*A*_2_, while with probability *p*_12_*/*(2*p*_2_) = (1 *− p*)(1 *− F*) they are *A*_1_*A*_2_. Therefore, the mean effect of the flip on the offspring’s phenotype satisfies

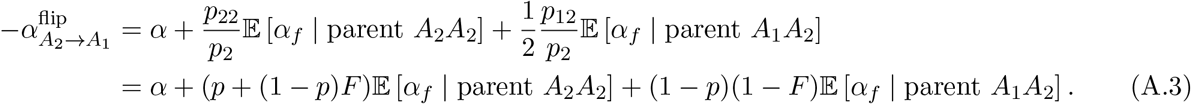

If we choose whether to flip *A*_1_ or *A*_2_ based on their frequencies *p*_1_ and *p*_2_, with the effect of the *A*_2_ *→ A*_1_ flip weighted negatively relative to that of the *A*_1_ *→ A*_2_ flip, we obtain a weighted average effect size of

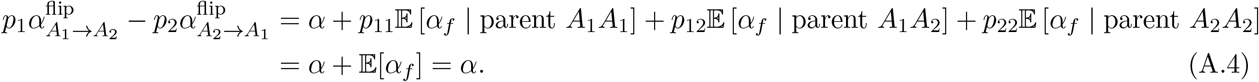

### A1.2 Estimates of the effect size of *A*_2_ produced by population and family-based association studies

#### Family-based association studies

We consider two family-based designs for an association study at the focal locus. In the sibling design, the difference between full siblings’ phenotypes is regressed on the difference in their genotypes. In the trio-based design, the offspring’s phenotype is regressed on their genotype, including as a covariate in the regression the average of their parents’ genotypes. These two designs return the same effect-size estimate in expectation, assuming no sibling indirect effects (we reproduce a proof of this below); we will generally find it more convenient to carry out our calculations for the sibling design.

#### Sibling design

Let *i* and *j* be siblings in family *f*, and define Δ*Y*_*f*_ = *Y*_*i*_ *− Y*_*j*_, Δ*g*_*f*_ = *g*_*i*_ *− g*_*j*_, and Δ*ϵ*_*f*_ = *ϵ*_*i*_ *− ϵ*_*j*_. A sibling association study returns an effect-size estimate

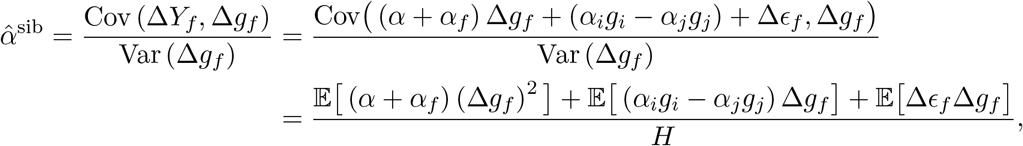

where *H* is the fraction of parents who are heterozygous at the focal locus. Since *α*_*i*_, *α*_*j*_, *ϵ*_*i*_, and *ϵ*_*j*_ are genotype-independent perturbations, 𝔼(*α*_*i*_*g*_*i*_ *− α*_*j*_*g*_*j*_) Δ*g*_*f*_ = 0 and 𝔼[Δ*ϵ*_*f*_ Δ*g*_*f*_] = 0, so

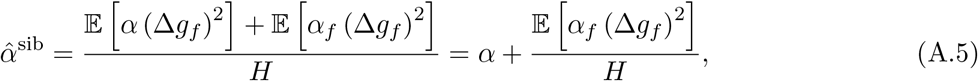

which deviates from *α* by an amount 𝔼*α*_*f*_ (Δ*g*_*f*_)^2^ */H*.

Let 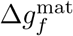 and 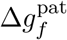 be the difference in the genotypes of siblings *i* and *j* in family *f* due to maternal and paternal transmission. 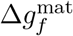 and 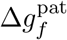 are linearly independent, and so the term additional to *α* in Eq. (A.5) can be split into 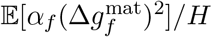 and 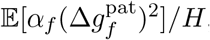, which we can analyze separately.

If the mother is heterozygous, then 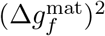 equals 1 with probability 1*/*2 and 0 with probability 1*/*2; if the mother is homozygous, then 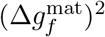 is 0. Therefore, denoting by *h*^m^ the event that the mother is heterozygous,

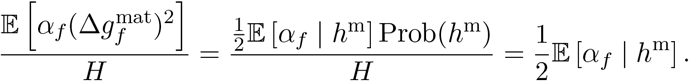

The same holds for paternal transmission, and so

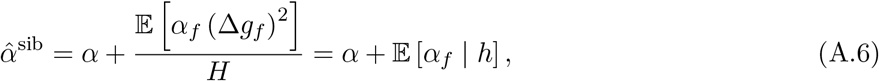

where *h* is the event that a chosen parent is heterozygous.

#### Trio design

We now show that this sibling-based estimate is, in expectation, the same as would be obtained from a trio-based design in which we regress offsprings’ phenotypes *Y* on their genotypes *g*, controlling for their midparents’ genotypes (*g*^*m*^ + *g*^*f*^)*/*2. Define the ‘segregation deviation’ *ς*_*i*_ of offspring *i* as the difference between their genotype and their midparent genotype:

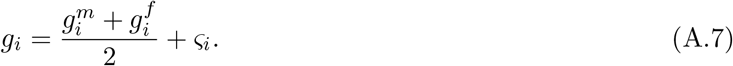

By the Frisch–Waugh–Lovell theorem (Angrist and Pischke 2009, pp. 35–36), the coefficient on the offspring’s genotype in the trio regression described above is the same as would be obtained if we (i) first regress offsprings’ genotypes on their midparents’ genotypes, and then (ii) regress offsprings’ phenotypes on the residuals from this first regression. However, the regression in (i) is Eq. (A.7), and so the residuals are just the *ς*_*i*_ values (in expectation). Therefore, the regression in (ii) yields a coefficient

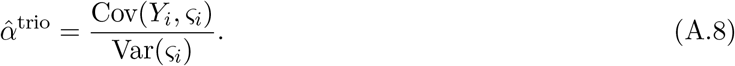

To show that this is equal, in expectation, to the sibling estimate, note that we can write the difference between two siblings’ genotypes as Δ*g*_*f*_ = *g*_*i*_ *− g*_*j*_ = *ς*_*i*_ *− ς*_*j*_. Moreover, *ς*_*i*_ and *ς*_*j*_ are linearly independent. Therefore, the sibling-based estimate can be written

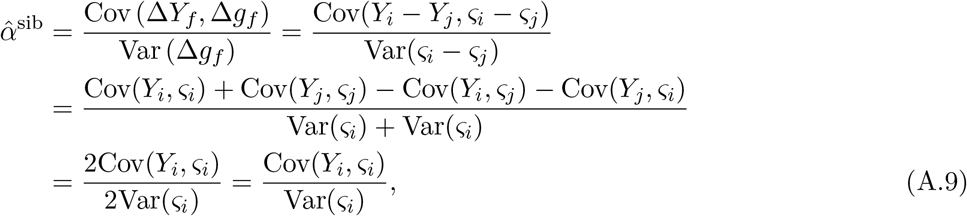

assuming Cov(*Y*_*i*_, *ς*_*j*_) = Cov(*Y*_*j*_, *ς*_*i*_) = 0, i.e., that there are no sibling indirect genetic effects. This is the same as the trio-based estimate in Eq. (A.8), and so the trio-based estimate is, in expectation, the same as in Eq. (A.6), i.e.,

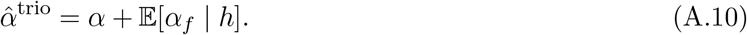

Therefore, assuming no sibling indirect effects, we can define a general ‘family-based’ estimate

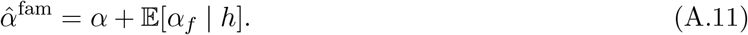

#### Population association study

Under the same one-locus model, with no environmental or genetic confounding, a population association study returns an effect-size estimate of

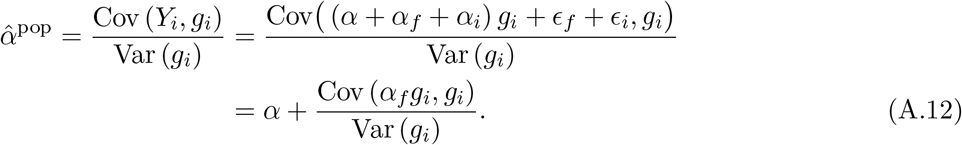

We can immediately see from Eq. (A.12) that if the family environments are randomized across genotypes, such that *α*_*f*_ and *g*_*i*_ are independent (implying Cov (*α*_*f*_ *g*_*i*_, *g*_*i*_) = 0), then the population estimate will coincide with *α*.

To calculate the deviation of the population estimate from *α* in the general case, let *F* be the inbreeding coefficient at the locus. Then Var(*g*_*i*_) = 2*p*(1 *−p*)(1 + *F*), where *p* is the frequency of the focal variant *A*_2_, and the frequency of heterozygotes is *p*_12_ = 2*p*(1 *−p*)(1 *−F*) while the frequencies of the two homozygotes are *p*_11_ = (1 *− p*)^2^ + *p*(1 *− p*)*F* (zero focal alleles) and *p*_22_ = *p*^2^ + *p*(1 *− p*)*F* (two focal alleles). The covariance term in Eq. (A.12) can then be written

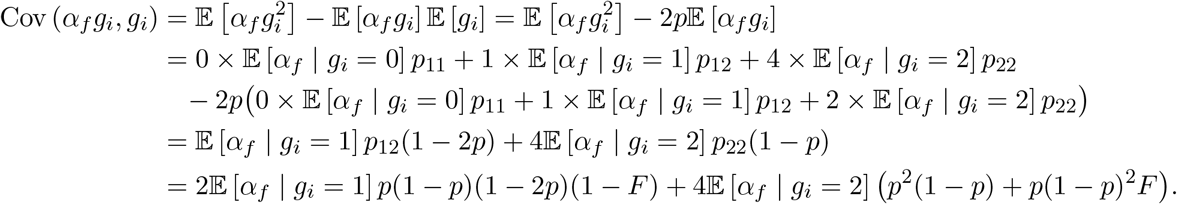

The population-based estimate is therefore

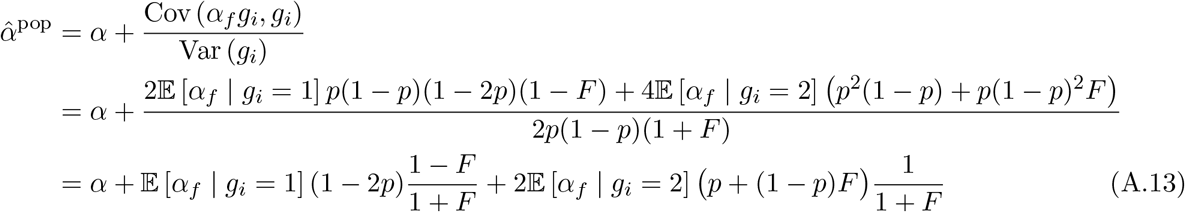

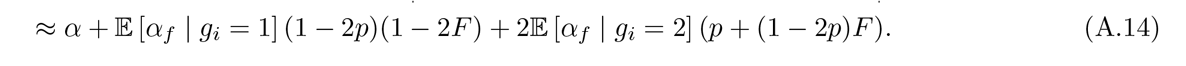

The approximation holds when *F* is small.

Using the fact that 𝔼 [*α*_*f*_] = 0, we can write Eq. (A.13) in terms of G*×*E effects conditional on the two homozygous offspring genotypes, 𝔼 [*α*_*f*_ | *g*_*i*_ = 0] and 𝔼 [*α*_*f*_ | *g*_*i*_ = 2]:

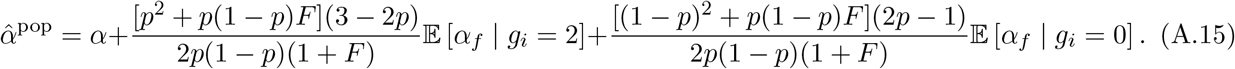

To facilitate comparison with Eqs. (6) and (2), it is useful to convert these offspring-genotype-conditioned G × E effects into parent-genotype-conditioned G×E effects. First, consider a 11 homozygous offspring. This individual inherited a 1 allele from their mother 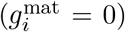, who could have been a 11 homozygote or a 12 heterozygote. For this offspring, we have

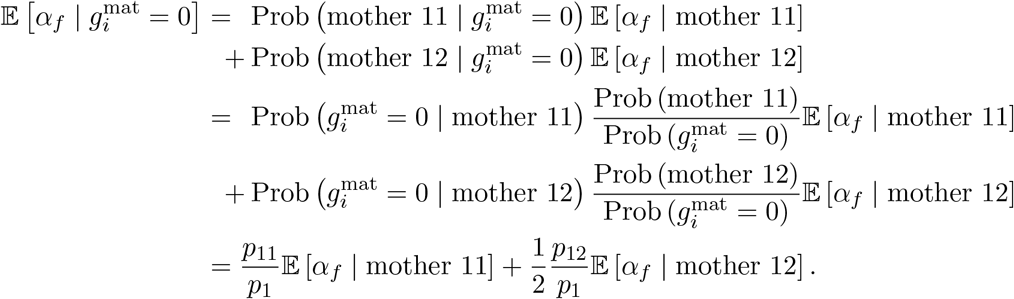

Similarly,

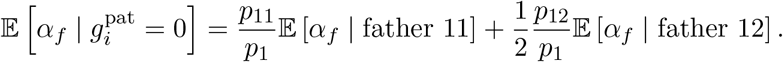

From these, and under the assumption that the inbreeding coefficient at the locus is not very large, it follows that

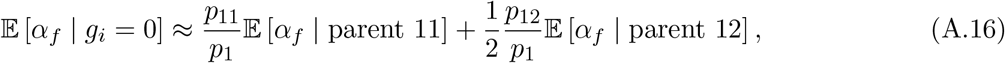

and, similarly,

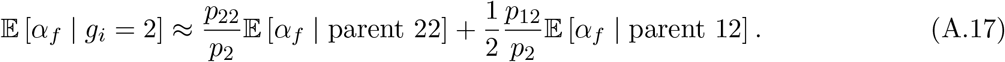

Substituting Eqs. (A.16) and (A.17) into Eq. (A.15), we obtain the population GWAS estimate, written in terms of G*×*E effects conditioned on the parental genotype:

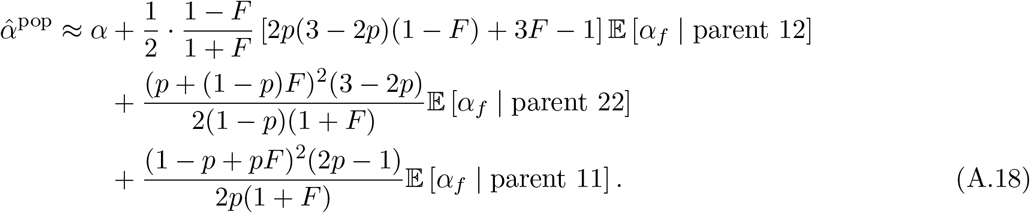

#### Writing the population-based estimate in terms of the family-based estimate

Again writing offspring *i*’s genotype in the form 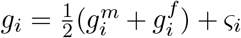 the population-based estimate can be written

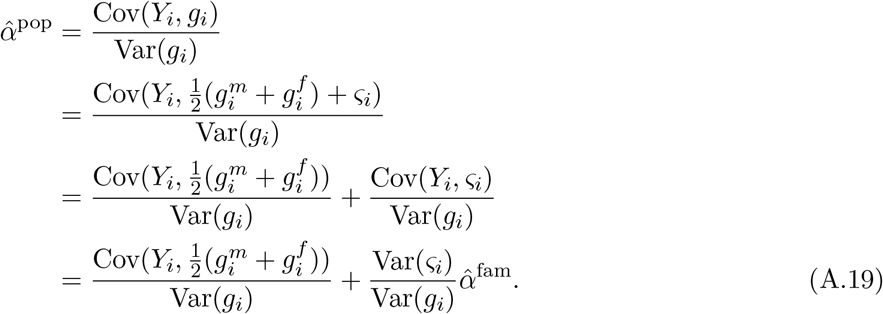

The first term is the slope based on variation across families, and the second is the slope based on variation within families due to the randomness of Mendelian segregation. Var(*g*_*i*_) = 2*p*(1 *− p*)(1 + *F*) is the genotypic variance at the locus, with *p* the frequency of *A*_2_ and *F* the inbreeding coefficient at the locus.

Write 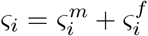, where 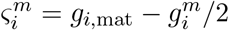 and 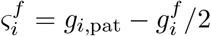 are genotypic deviations due to segregation in the mother and the father respectively, taking on the value 0 if the parent is homozygous at the locus and the values *−*1*/*2 or 1*/*2 (each equally likely) if the parent is heterozygous. Note that 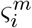 and 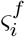 are linearly independent and have mean zero. Therefore,

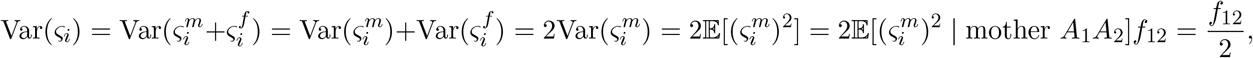

where *f*_12_ = 2*p*(1*−p*)(1*−F*) is the fraction of parents who are heterozygous at the locus (genotype *A*_1_*A*_2_). Substituting this into Eq. (A.19),

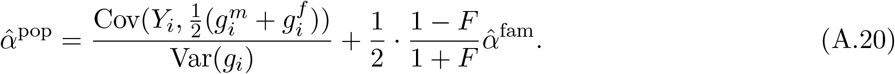

As shown above,

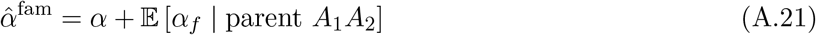

and

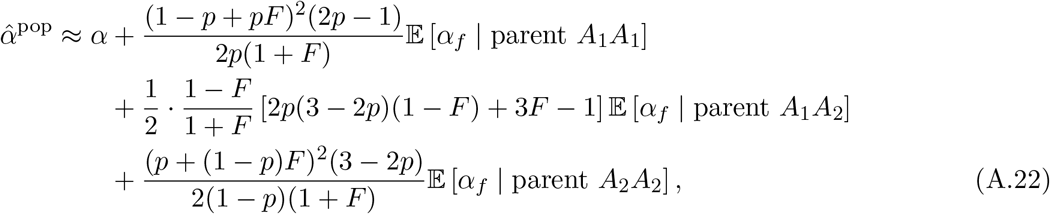

so

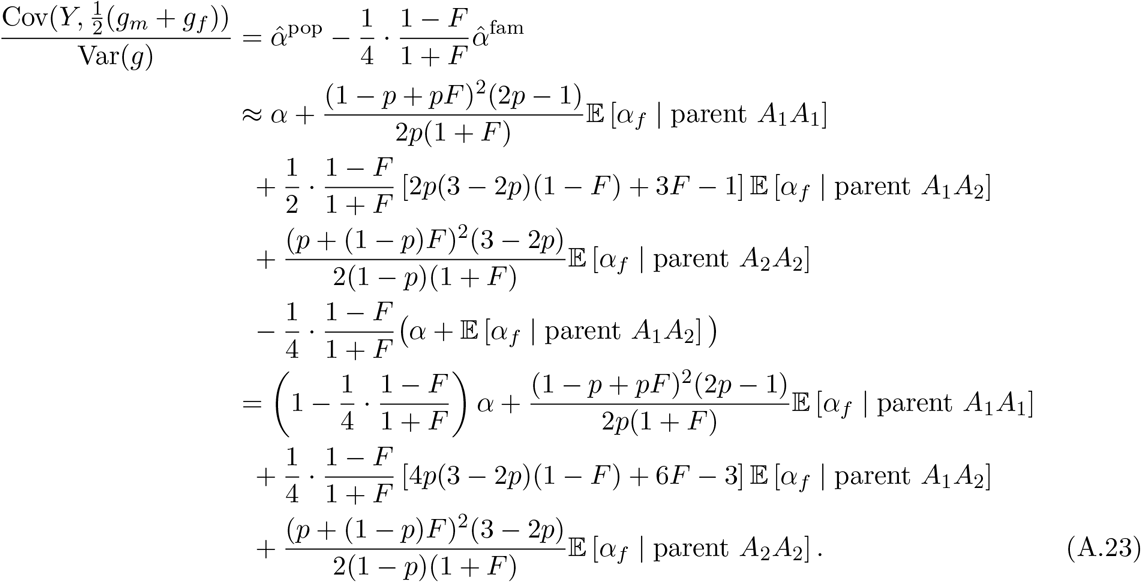

### A1.3 Relationship to the ‘average effect’ defined in Lee & Chow (2013)

Lee and Chow (2013) propose a definition of the ‘average effect’ of an allele at a biallelic locus when the effect of the allele depends on the environment. Again, let the alleles at the locus be *A*_1_ and *A*_2_, and let *P*, 2*Q*, and *R* be the frequencies of the genotypes *A*_1_*A*_1_, *A*_1_*A*_2_, and *A*_2_*A*_2_ respectively. Lee and Chow (2013) assign the label *ε*_*i*_ to environment *i*, and in their Eq. (15) define the average effect of allele *A*_2_ as

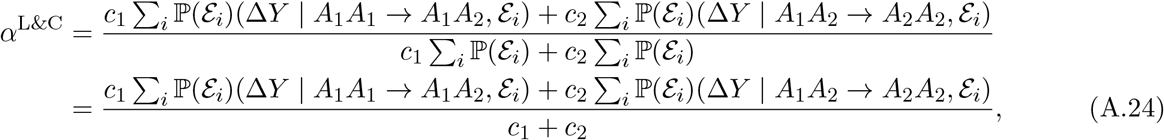

where *c*_1_ = *P* (*Q* + *R*), *c*_2_ = *R*(*P* + *Q*), ℙ(*ε*_*i*_) is the frequency of environment *ε*_*i*_, and (Δ*Y* | *g → g*^*′*^, *ε*_*i*_) is the mean phenotypic effect of experimentally switching genotype *g* to genotype *g*^*′*^in environment ε_*i*_.

To relate this to the parameters in our phenotypic model *Y* = *Y* ^*\**^ + (*α* + *α*_*f*_)*g* + *ϵ*, note that we can simply consider the environment of each family *f ∈ F* as a distinct environment labelled *f* . Then ℙ (*f*) = 1*/*|*F*| and

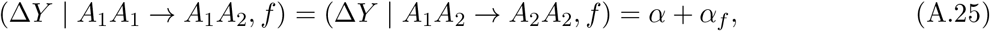

with the latter equality relying on the assumption of additivity (i.e., no dominance) at the locus. Substituting these values into Lee & Chow’s definition of the average effect of *A*_2_,

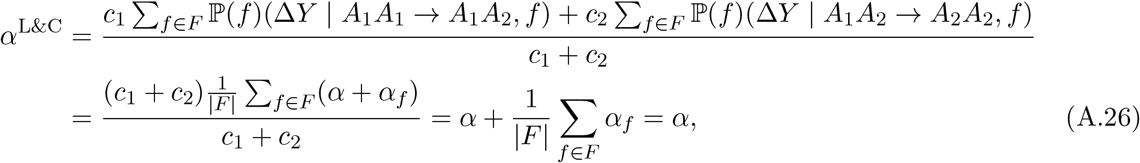

since 𝔼[*α*_*f*_] = 0, by assumption.

### A1.4 Main Text example 1

As we have seen, in the presence of G*×*E interactions, family- and population-based association studies at a causal locus can return different estimates from each other and from the true causal effect when genotypes are nonrandomly distributed across interacting environments. As a simple example that illustrates this point, consider a G*×*E model in which there are two populations. In the environment inhabited by the red population, the effect of allele *A*_2_ is *α* + *x*, while in the environment inhabited by the blue population, the allele’s effect is *α − x*. So, with reference to our phenotypic model (A.1), *α*_*f*_ = +*x* in the red population and *α*_*f*_ = *−x* in the blue population. Children are raised in the environment of their parents. The frequency of *A*_2_ is *p*_1_ in the red population and *p*_2_ in the blue population, and each population is in Hardy-Weinberg equilibrium at the locus.

The mean trait values are then 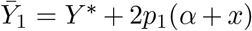 in the red population and 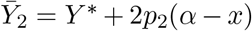 in the blue population. Because effects are not heterogeneous within each population, unconfounded population and family-based association studies *within* each population would return effect-size estimates of *α* + *x* in the red population and *α−x* in the blue population. If we were to draw a large sample equally from both populations, the average effect of the allele *A*_2_ that would be obtained by the manipulation-based approach from the sample is *α*.

A population-based association study across the same sample would return, in expectation, an effect-size estimate 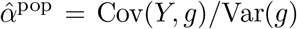. Denote by 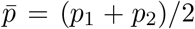 the frequency of *A*_2_ in the sample and by 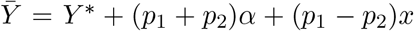 the average trait value in the sample. From the law of total covariance, using the subscript notation ‘pops’ to denote that an expectation, variance, or covariance is taken across populations and ‘pop’ to denote conditioning on a particular population,

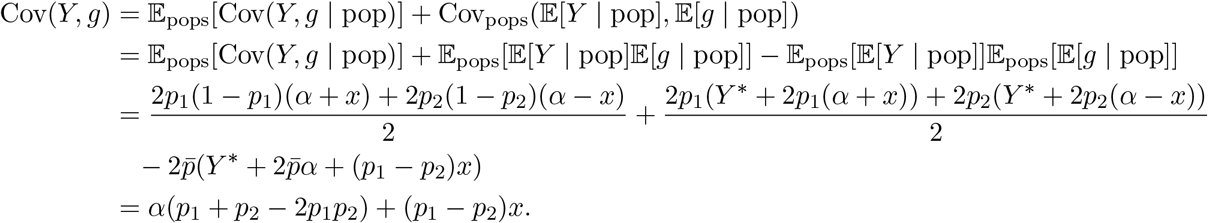

From the law of total variance,

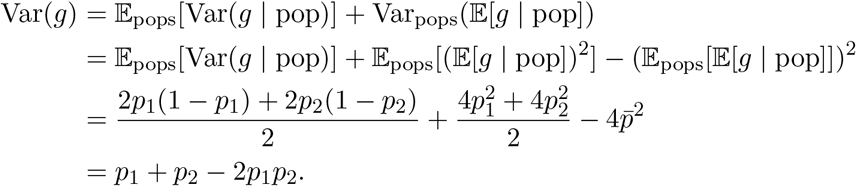

Therefore,

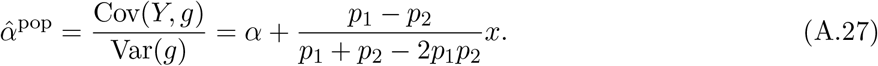

That is, the deviation from *α* of the effect-size estimate of a population-based association study is driven by the difference in allele frequencies between the two populations, *p*_1_ *− p*_2_.

Using Eq. (A.10), and recalling the notation *h* for the event that a chosen individual is heterozygous at the focal locus, a population-based association study across the sample returns an effect-size estimate

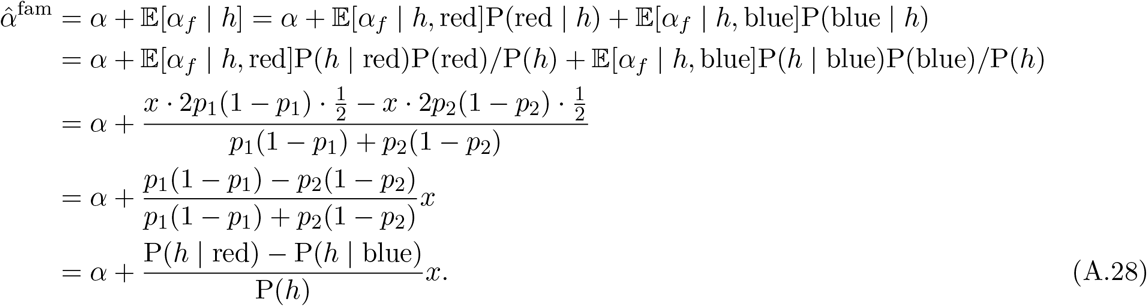

That is, the deviation from *α* of the effect-size estimate of a family-based study is driven by the difference in the heterozygosities of the two populations, 2*p*_1_(1 *− p*_1_) *−* 2*p*_2_(1 *− p*_2_).

#### Simulation details

For Fig. 1 in the Main Text, we simulate population-based and family-based GWASs under this model. To allow allele frequencies to vary between the populations, we simulate the two populations having experienced drift in allele frequencies away from an ancestral frequency. At each SNP, we simulate independent drift in each population under a beta-binomial approximation (Balding and Nichols 1995; Balding 2003), from an ancestral frequency of 0.5 and with an *F* = 0.05. To model phenotypes we set *α* = 1 and *x* = 0.1 and we add a small amount of environmental noise with a standard deviation of 0.1. We sample 10,000 diploid individuals in each population and conduct a population-based GWAS in the combined sample. We calculate the effect-size estimate from the family-based association study as the average of the two effect sizes weighted by fraction of heterozygotes in each population.

### A1.5 Linkage disequilibrium between marker and causal loci

There is a causal locus segregating for alleles *A* and *a* and a marker locus segregating for alleles *M* and *m*. The effect of *A* at the causal locus is *α*, independent of the environment (no G*×*E). For simplicity, we assume Hardy-Weinberg genotype frequencies at the marker locus.

First, we are interested in the average phenotypic effect of randomly choosing an *m* allele in a zygote and flipping its haplotype to a random one containing *M* . If the flipped haplotype does not contain the causal locus, the flip obviously has no phenotypic consequences. If the flipped haplotype does contain the causal locus, then the probability that there is an *a→A* flip at the causal locus is *p*_*a*|*m*_*p*_*A*|*M*_, while the probability that there is an *A→a* flip at the causal locus is *p*_*A*|*m*_*p*_*a*|*M*_, where we use the notation *p*_*x*|*y*_ for the proportion of *y*-carrying haplotypes that also carry allele *x*. The average effect of the *m→M* flip is therefore

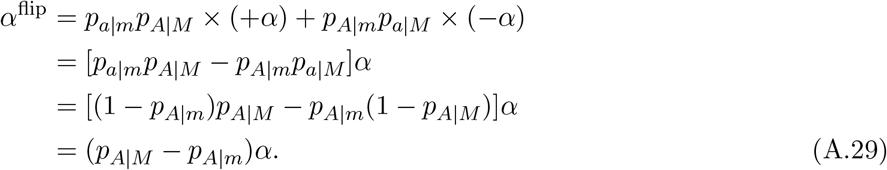

To calculate the effect size estimated by a population-based association study at the marker locus, we calculate the coefficient of LD *D* between the marker and causal loci, in terms of the quantities *p*_*x*|*y*_, and substitute these into the standard GWAS effect size formula 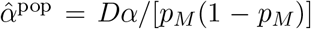 (assuming Hardy-Weinberg genotype frequencies at the marker locus). We have

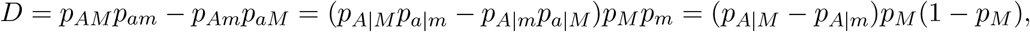

where the *p*_*xy*_ and *p*_*y*_ are haplotype and genotype frequencies, respectively. So

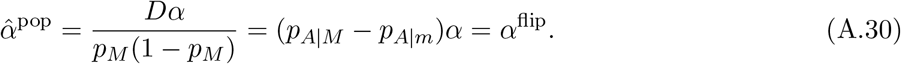

To calculate the effect size estimated by a family-based association study at the marker locus, we consider a sibling design (as noted in Section A1.2 above, the trio-based design will return the same estimate in expectation, assuming no sibling indirect effects). Let *g*_*ℳ*_ be the number of copies of allele *M* carried by an offspring at the marker locus, with 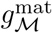 and 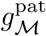 the number of copies inherited maternally and paternally, respectively. Let *g*_*𝒜*_, 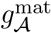, and 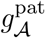 be the analogous quantities for allele *A* at the causal locus. As in Section A1.2, we denote by Δ a difference between two full siblings for some quantity (genotype or phenotype). Finally, we denote by *h* the event that a parent is heterozygous at the marker locus, with *H* the associated frequency of heterozygotes, and we denote by *h*^c^ the event that a parent is a coupling double-heterozygote (genotype *AM/am*) and by *h*^r^ the event that a parent is a repulsion double-heterozygote (genotype *Am/aM*), with *H*^c^ and *H*^r^ the associated frequencies (we assume genotype frequencies to be the same for mothers and fathers).

Given this notation, a sibling association study at the marker locus returns an estimate

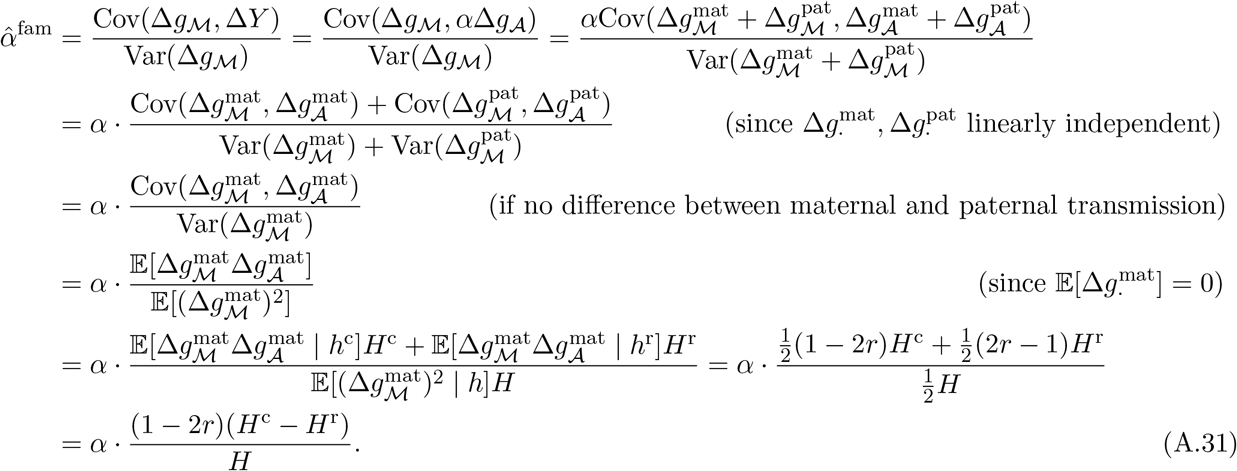

By the notation used in the Main Text, *p*′_*A*/*M,Mm*_ and *p*′_*A*/*M,Mm*_ are the proportions in *Mm* individuals of *M* - and *m*-containing haplotypes, respectively, that also contain *A*. We have

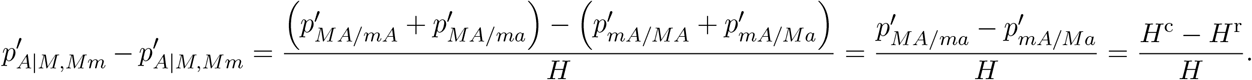

Therefore,

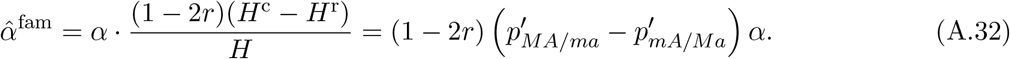

If the marker and causal loci are tightly linked (*r ≈* 0) and genotype frequencies are approximately constant in the parent and offspring generations (*p*^*′*^_*MA/ma*_ *≈ p*_*MA/ma*_ and *p*^*′*^_*mA/Ma*_ *≈ p*_*mA/Ma*_), then

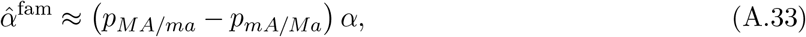

which is Eq. (10) in the Main Text.

#### Decomposing the total LD into contributions from marker-homozygotes and heterozygotes

Our aim is to decompose the overall degree of LD between the alleles *M* and *A* in the sample, *D*, into a contribution from LD among individuals heterozygous at the marker locus, *D*_het_, and a contribution from LD among individuals homozygous at the marker locus, *D*_hom_.

To do so, we first consider the more general case of two disjoint subpopulations 1 and 2 in which the frequency of *M* is *p*_*M* |1_ and *p*_*M* |2_ respectively, the frequency of *A* is *p*_*A*|1_ and *p*_*A*|2_ respectively, and the degree of LD between *M* and *A* is *D*_1_ and *D*_2_ respectively. In a sample that weights populations 1 and 2 by *f*_1_ and *f*_2_ = 1 *− f*_1_, the degree of LD between *M* and *A* is

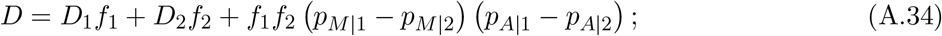

i.e., the weighted average of the LDs between the two subpopulations plus a contribution to overall LD from allele frequency differences between the subpopulations at both loci (Nei and Li 1973).

In our case, population 1 comprises all individuals in the sample who are heterozygous at the marker locus (genotype *Mm*) while population 2 comprises all individuals who are homozygous at the marker locus (genotypes *mm* and *MM*). Noting that *p*_*M* |het_ = 1*/*2 by definition, we have, from Eq. (A.34),

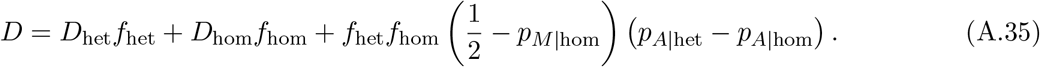

We can also calculate *D*_het_ and *D*_hom_ explicitly. If we randomly pick a haplotype, let the random variable *X* take the value 1 if the haplotype carries *M* and 0 if it carries *m*, and let *Y* take the value 1 if the haplotype carries *A* and 0 if it carries *a*. The degree of LD between the loci in the group of individuals heterozygous at the marker locus is

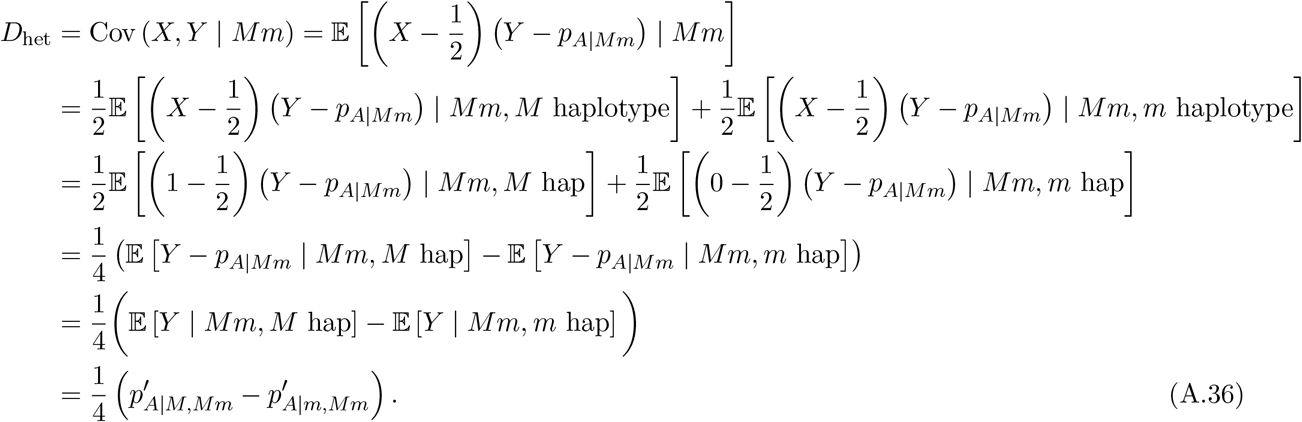

The degree of LD between the loci in the group of individuals homozygous at the marker locus is

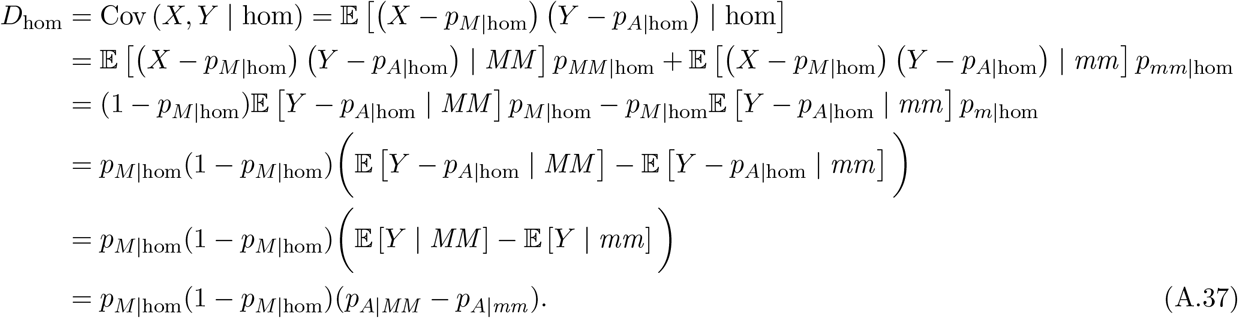

For comparison, note that we can write the overall LD as *D* = *p*_*M*_ (1 *− p*_*M*_)(*p*_*A*|*M*_ *− p*_*A*|*m*_).

### A1.6 Main Text example 2

As a simple illustration of the effects of LD between marker and causal loci, suppose again that there are two populations, 1 and 2. The frequencies of allele *A* at the causal locus and allele *M* at the marker locus are *p*_*A*|1_ and *p*_*M* |1_ in population 1 and *p*_*A*|2_ and *p*_*M* |2_ in population 2, respectively. The degrees of LD between *M* and *A* in populations 1 and 2 are *D*_1_ and *D*_2_. We allow these allele frequencies and strengths of LD to differ between the two populations. For simplicity, we assume that mating is random in the two populations so that Hardy-Weinberg equilibrium prevails within each.

The expected frequencies of alleles *M* and *A* in a sample that weights the two populations equally are *p*_*M*_ = (*p*_*M* |1_ + *p*_*M* |2_)*/*2 and *p*_*A*_ = (*p*_*A*|1_ + *p*_*A*|2_)*/*2. The coefficient of (cis-)LD between *M* and *A* in the sample is

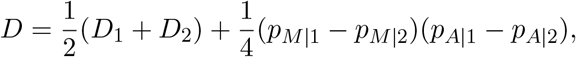

with contributions from LD within the two populations, (*D*_1_ +*D*_2_)*/*2, and an additional contribution from allele frequencies difference between them at the two loci, (*p*_*M* |1_ *− p*_*M* |2_)(*p*_*A*|1_ *− p*_*A*|2_)*/*4 (cf. Nei and Li 1973). The coefficient of trans-LD between *M* and *A* in the sample, caused by their frequency differences between the two populations, is 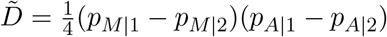.

An unconfounded association study at the marker locus, carried out separately in populations 1 and 2, would return effect sizes 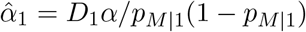 and 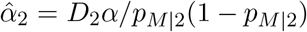 for allele *M* . These correspond to the haplotype-flipping effect sizes of the marker allele *M* in the two populations, as described above.

A population-based association study at the marker locus across a sample that weights the two populations equally returns an effect-size estimate

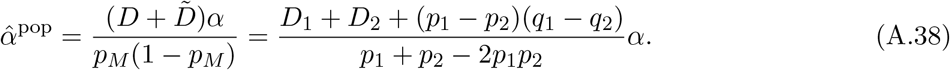

The numerator is (twice) the total population-wide degree of LD between *M* and *A*, with contributions from LD within the two populations, *D*_1_ + *D*_2_, and an additional contribution from allele frequencies difference between them at the two loci, (*p*_1_ *− p*_2_)(*q*_1_ *− q*_2_) (cf. Nei and Li 1973).

On the other hand, the family-based GWAS effect-size estimate is

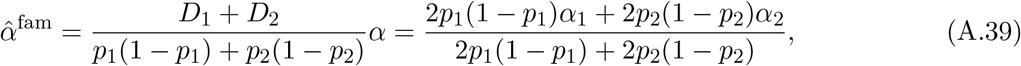

i.e., the heterozygosity-weighted average of the population-specific effects.

## A2 Defining the causal effect of the PGS in terms of allele-flipping manipulations

As described in the Main Text, the scenario we are interested in with respect to PGSs is one where an investigator has access to effect-size estimates 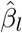 at a set of genotyped loci *l ∈* Λ from a prior GWAS on trait X, and uses these to construct a trait-X PGS for each individual *i* in a sample that does not overlap with the original GWAS sample:

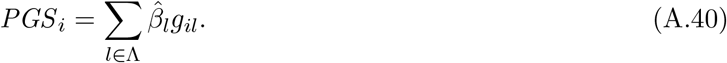

The value of trait Y (which could be the same as trait X, or not) for individual *i* in family *f* follows the linear model

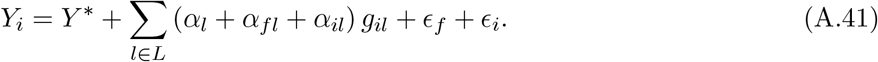

For simplicity, we have assumed that the PGS loci, Λ, are a subset of the loci that causally affect trait Y, *L*, and that the loci in *L* are in Hardy-Weinberg and linkage equilibrium.

We first consider the average effects on the value of trait Y of hypothetical experimental manipulations of genotypes at the loci in Λ.

### A2.1 Single-locus manipulation

Initially, we consider the effect of ‘flipping’ (as if by CRISPR) a single allele at one of the PGS loci in a gamete (before it fuses to form a zygote), and measure the slope of the average shift in the phenotype caused by this flip against the change in PGS. Given that there are many PGS loci, all with different effect-size estimates and allele frequencies, the question is how to choose the locus at which the allele flip will occur. We want a manipulation that is biologically relevant, and so we want to preferentially flip alleles at loci that are common sources of PGS variation among our sampled individuals. A natural choice is to choose loci with probability proportional to their contribution to the PGS sample variance; that is, locus *l* is chosen with probability 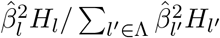 (note that this is the PGS variance rather than the additive genetic variance contributed by PGS loci, because the investigator does not have access to the true effect sizes of alleles at the loci, only estimates of these effect sizes from a prior GWAS).

Having chosen a locus, we randomly sample a gamete. If it carries an *A*_1_ allele at the chosen locus, we flip this to *A*_2_, while if it contains an *A*_2_ allele, we flip it to *A*_1_. The changes in the PGS from these two flips are 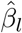 and 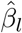 respectively. If we average the flip over all individuals in our sample, we would observe an average phenotype to PGS slope of 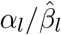 (note that the family and individual G*×*E effects drop out here, since they are zero on average—see discussion around Eq. (4)).

Given the weightings with which we choose the various loci in Λ, the expected effect of this allele-flipping scheme (proportionate to the change in the PGS) is therefore

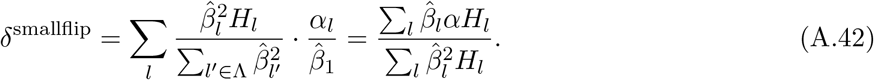

Note that while this represents a reasonable definition of the effect, different choices of how we pick loci to flip would result in different *δ*s.

### A2.2 Multi-locus manipulations

We can also define hypothetical experimental manipulations in which alleles are flipped at many PGS loci.

The investigator chooses a random pair of zygotes A & B from the sample, and manipulates, by allele flipping, the genotype of A so that it matches that of B at the PGS loci. The investigator does not alter A’s genotype at other loci, and A grows up in the same environment that they would have in the absence of the genetic manipulation. The manipulation changes A’s PGS from *PGS* _*A*_ to *PGS* _*B*_. The investigator is interested in the value of A’s phenotype given this manipulation.

Let *g*_*A,l*_ and *g*_*B,l*_ be A’s and B’s genotype at locus *l*. Denote by 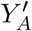 the phenotypic value of A if the investigator performs the manipulation, and by *Y*_*A*_ the phenotypic value if they do not.

The investigator performs this manipulation over a large number of random pairs of individuals. We consider two measurements (1 & 2 below) that the investigator might be interested in, which, as we show, yield the same value in the absence of G*×*E but differ in the presence of G*×*E. Measurement 1 results in the same effect as our randomization and single locus PGS manipulation. Measurement 2 results in a different quantity, highlighting how relatively minor decisions can result in quite different quantities being dervied.

#### Measurement 1

The investigator regresses the counterfactual phenotypic values 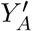 on the individuals’ new polygenic scores *PGS* _*B*_, yielding an expected coefficient

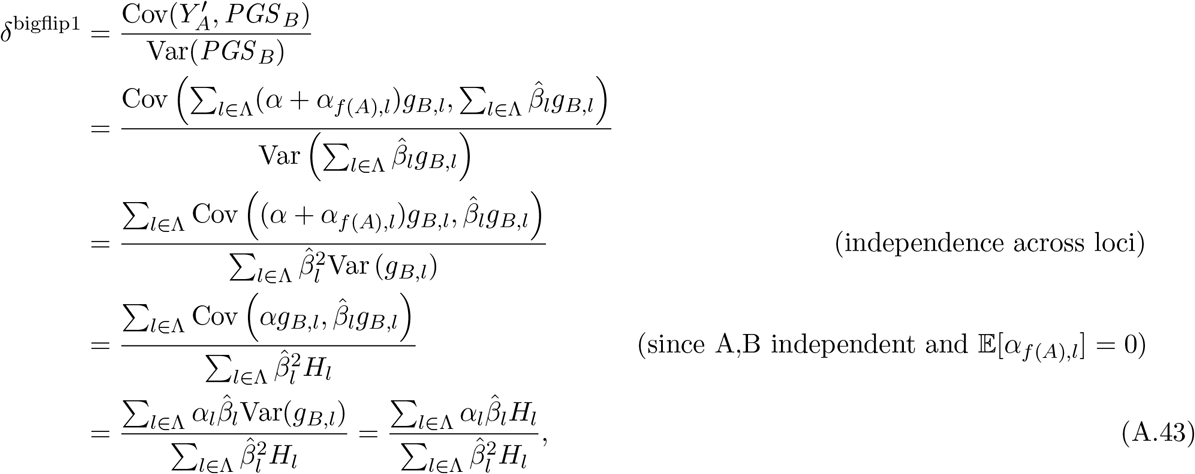

which is the same as Eq. (A.42) and the ‘randomization’ value in the Main Text (Eq. (14)) that would obtain if we randomly assigned PGS genotypes independent of environmental and genetic backgrounds . The reason is that, in a sense, the manipulation is testing the effect of B’s PGS in a random genetic and environmental background (since A is chosen randomly from the sample).

#### Measurement 2

The second measurement the investigator might be interested in conducting is to regress the change in the phenotype caused by the manipulation, 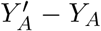, on the change in the PGS, *PGS* _*B*_ *− PGS* _*A*_. This yields, in expectation,

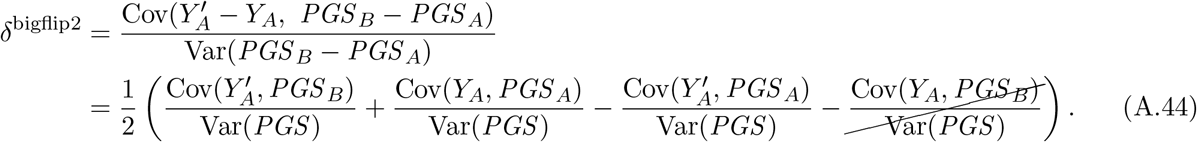

As before, 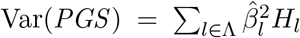. The first term in brackets is *δ*^bigflip1^. The second term is the population slope of the PGS, *δ*^pop^. Expanding the covariance in the third term:

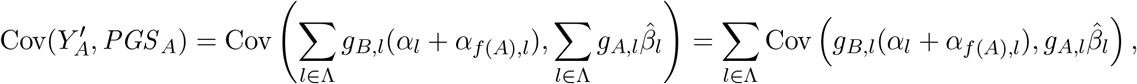

assuming no LD among loci. Consider the summand, letting 2*p*_*l*_ = 𝔼 [*g*_*A,l*_] = 𝔼 [*g*_*B,l*_]:

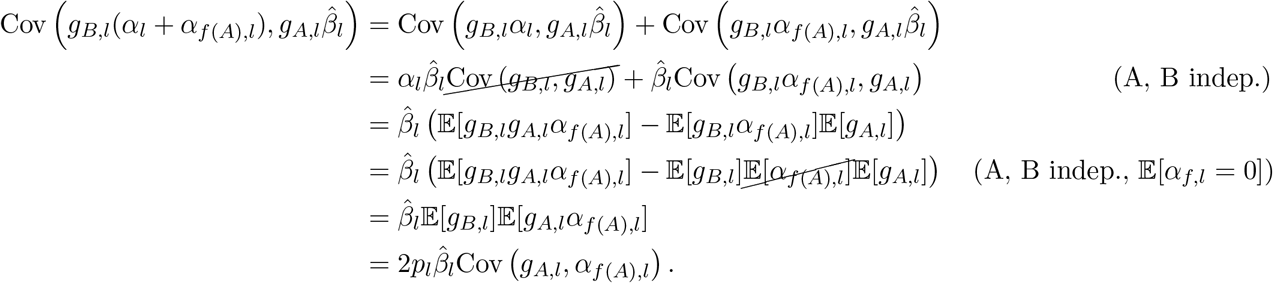

Putting all of this together,

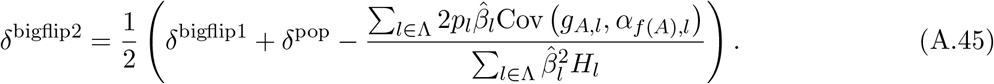

Since

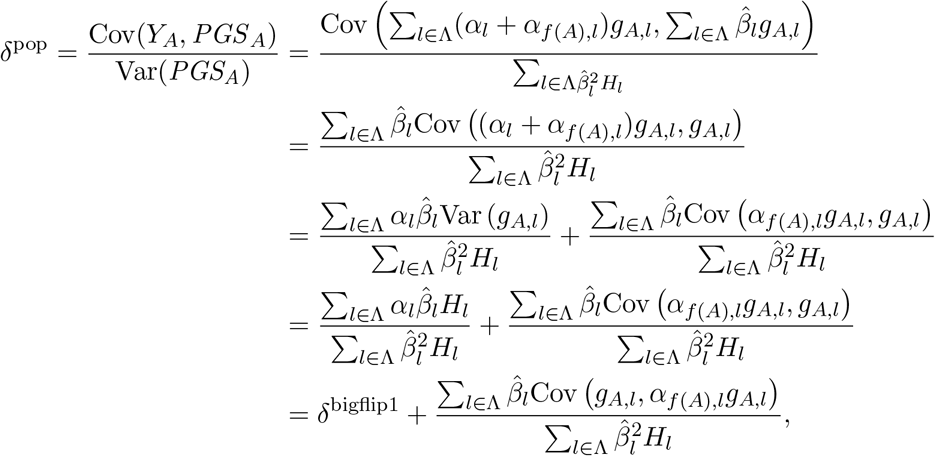

*δ*^bigflip2^ can also be written

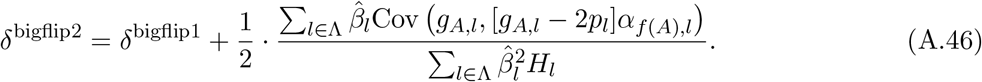

Clearly *δ*^bigflip1^ and *δ*^bigflip2^ need not agree in general when there is G*×*E and PGS genotypes are not homogeneously distributed across environments. The key difference between the two measurements is that when we take the difference between pairs we are seeing the effect of swapping the genotype of A to that of B in A’s environment—environments are therefore not randomized in this measurement.

### A3 Family-based PGS regression

Given a PGS for trait X,

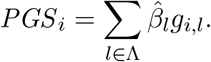

we are interested in its relationship with the value of trait Y (which could be the same as X, or not). Again, the value of trait Y follows the linear model

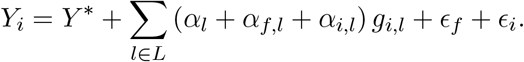

We are specifically interested in the slope estimate 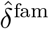 produced by least-squares estimation of the regression

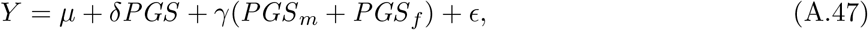

where *PGS* is the PGS of the offspring and *PGS* _*m*_ and *PGS* _*f*_ are the PGSs of the mother and father, respectively.

#### A3.1 Equivalent estimates of *δ* from trio- and sibling-based designs

The estimate of *δ* obtained from the regression in Eq. (A.47) is the same, in expectation, as would be obtained from the regression of siblings’ phenotypic differences on the differences in their PGSs. To see this, write the PGS of individual *i* as

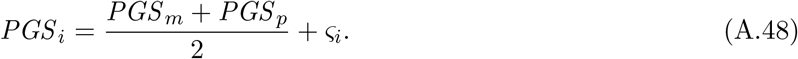

*ς*_*i*_ is the deviation of *i*’s PGS from the midparent value due to the randomness of parental transmission.

From the Frisch–Waugh–Lovell theorem (Angrist and Pischke 2009, pp. 35–36), the estimate 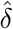 obtained by least-squares estimation of the multivariate regression (A.47) is the same as would be obtained if we first regressed *PGS* _*i*_ on *PGS* _*m*_ +*PGS* _*p*_, extracted the residuals from this first-stage regression, and regressed *Y*_*i*_ on the residuals. Since the the first-stage regression is simply Eq. (A.48), the residuals are, in expectation, the *ς*_*i*_, and so the estimate of *δ* in the second-stage regression is

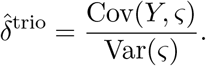

Now suppose that we have pairs of full siblings *i* and *j*, and regress the difference in their phenotypes, *Y*_*i*_ *− Y*_*j*_, on the differences in their PGSs, *PGS* _*i*_ *− PGS* _*j*_ = *ς*_*i*_ *− ς*_*j*_. This yields an estimate of *δ* of

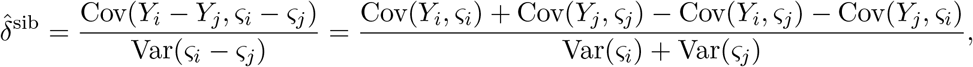

since *ς*_*i*_ and *ς*_*j*_ are linearly independent. Assuming no indirect effects of sibling *i*’s genotype on sibling *j*’s phenotype, and vice versa, Cov(*Y*_*i*_, *ς*_*j*_) = Cov(*Y*_*j*_, *ς*_*i*_) = 0, and so

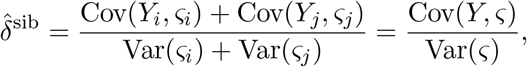

the same as the trio-based estimate, in expectation.

#### A3.2 Calculating the family-based estimate of *δ*

To calculate 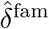, we make use of the equivalence (in expectation) of the estimates produced by the trio and sibling designs, and focus for simplicity on the sibling design, which produces an estimate of the form

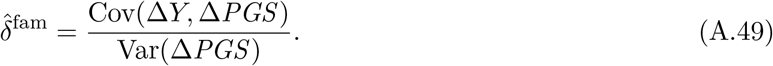

The covariance between siblings’ differences in their PGSs and differences in their trait values is

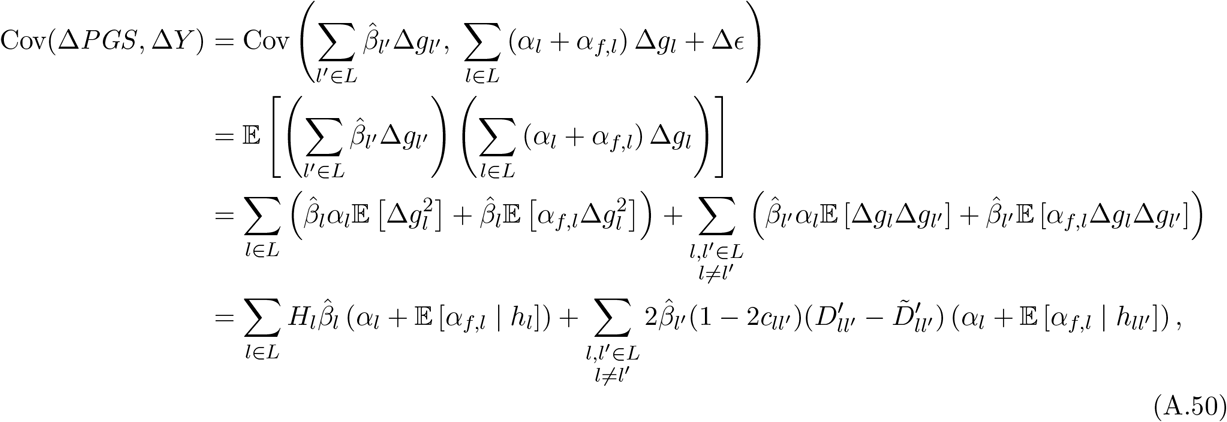

where *h*_*l*_ is the event that a parent is heterozygous at *l* (with associated probability *H*_*l*_), 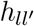 is the event that a parent is doubly heterozygous at *l* and *l*^*′*^, 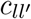 is the recombination rate between *l* and *l*^*′*^, and 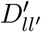 and 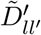 are the degrees of cis- and trans-LD, respectively, between loci *l* and *l*^*′*^ in parents (for similar calculations and more on the sources of cis and trans LD, see Veller and Coop 2023). The calculation relies on the assumption that the conditional expectation of *α*_*f,l*_ is the same for parents who are coupling and repulsion double heterozygotes, which might be a problematic assumption, e.g., in admixed populations.

The variance of the difference in siblings’ PGSs is

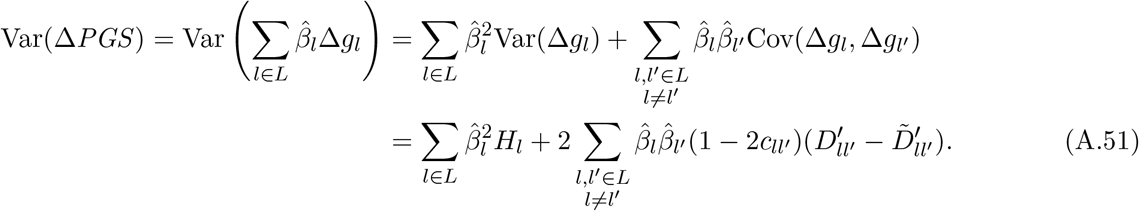

If the cis- and trans-LD terms are equal (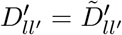, e.g., due to there being no genetic confounding— for more details, see Veller and Coop 2023), Eq. (A.50) simplifies to

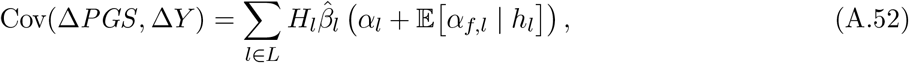

while Eq. (A.51) simplifies to

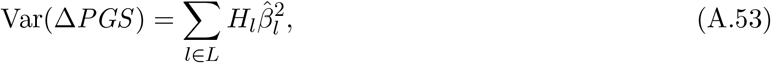

so that the estimate of *δ* produced by the sibling design (A.49), and therefore also by the trio design (in expectation), is

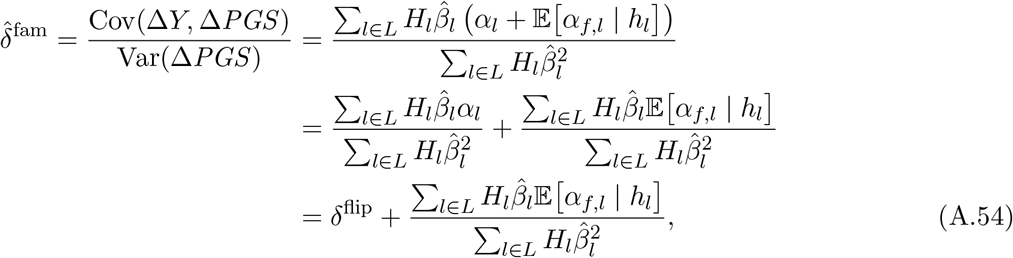

where we have labelled *δ*^flip^ = *δ*^smallflip^ = *δ*^bigflip1^ for the manipulation-based definitions of the effect of the PGS defined above (which also equal the randomization definition *δ*^rand^ defined in the Main Text).

Therefore, 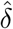 will be biased away from the mainpulation/randomization definitions of the effect of the PGS to the extent that heterozygous parents are distributed nonrandomly across interacting environments and therefore potentially exhibit unusual values of *α*_*f,l*_ (recall that, across the population as a whole, 𝔼*α*_*f,l*_ = 0).

The underlying reason is that the within-family variation in *PGS* on which the estimate Eq. (A.49) (and also the trio-based estimate) is based comes only from loci at which parents are heterozygous; therefore, the resulting estimate of *δ* ‘sees’ only the G*×*E effects that apply to heterozygous parents. In contrast, the manipulation definitions of the effect *δ* of the PGS involve flipping alleles that were inherited from both homozygous and heterozygous parents, and, similarly, the randomization definition involves placing individuals in environments chosen from the full population distribution, i.e., including those environments inhabited by homozygous parents. Thus, if heterozygous parents tend to have systematically different environments to homozygotes, the effects we estimate from families will not agree with the allele-flipping and manipulation definitions of the effect of the PGS.

#### A3.3 Is there an interpretation of the within-family PGS slope as being a weighted sum across families?

One scenario where a causal interpretation of 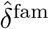 as an average of family-specific slopes *δ*_*f*_ can potentially be rescued is where each family has a family-specific G*×*E multiplier that commonly influences effect sizes at all loci. That is, each family *f* has a value *C*_*f*_ such that 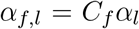 for all *l ∈ L*. Within family *f*, then,

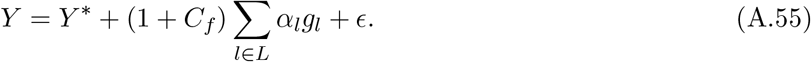

This corresponds to a scenario where some families occupy environments that amplify the effects of alleles on the trait (*C*_*f*_ *>* 0) while others occupy environments that attenuate allelic effects (*C*_*f*_ *<* 0). A similar scenario has been proposed for sex differences in the effects of alleles on various traits (Zhu et al. 2023). For simplicity, we assume that the PGS is constructed for the same trait Y. Assume further that the effect-size estimates used to construct the PGS, 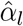, are inflated from the true effect sizes *α*_*l*_ by a multiplier *B* that is constant across loci (allowing also for uncorrelated noise). That is, 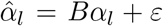, with the zero-mean term *ε* uncorrelated with *α*_*l*_ (and with the *p*_*l*_, etc.). Note that this nests the special case where the effect-size estimates are unbiased (*B* = 1).

Under these conditions, each family is characterized by a PGS slope *δ*_*f*_ = (1 + *C*_*f*_)*/B*, such that any genetic manipulation that increases the PGS of an offspring in family *f* by 1 unit—no matter which loci are flipped and in what direction—increases the offspring’s trait value by *δ*_*f*_ in expectation. Our aim here is to determine if the regression coefficient 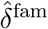 can be interpreted as an average of these family-specific slopes *δ*_*f*_ .

Let the set of families in the sample be *F* . We define the indicator 𝟙_*l,f*_ to take the value 1 if a randomly chosen parent in family *f* is heterozygous at locus *l* and 0 otherwise. Then we can write

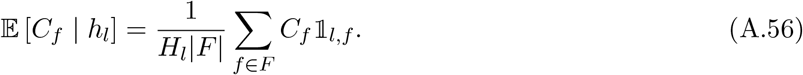

Eq. (A.54) then becomes

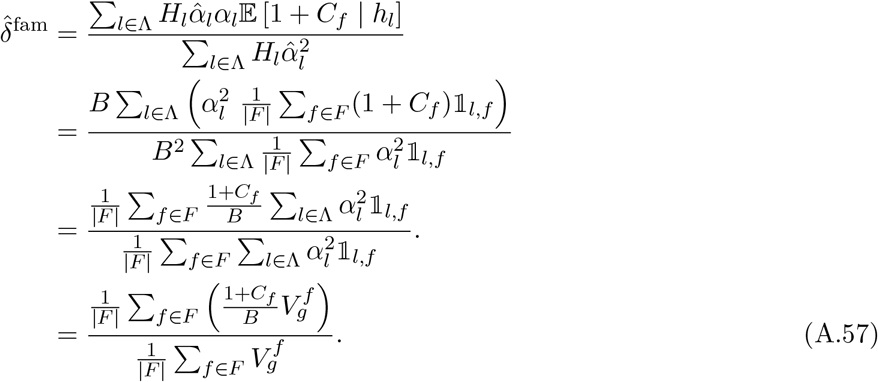

This is a weighted average across families of the family specific multipliers *δ*_*f*_ = (1 + *C*_*f*_)*/B*. The weight of family *f* in this average, 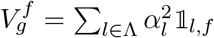, is the segregation variance for the PGS in family *f*, with the value higher if the parents in family *f* are heterozygous at more of the loci in Λ.

Note that the unweighted average of *C*_*f*_ is zero, since 0 = 𝔼 *α*_*f,l*_ = *α*_*l*_𝔼 [*C*_*f*_], and so, in general, 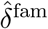 will be a biased estimate of the population-average value of *δ*_*f*_ if families in which parents are more heterozygous at PGS loci, and which therefore exhibit greater segregation variance, are associated with unusual values of *C*_*f*_ .

This is a particular instance of a more general phenomenon regarding least-squares regression, which has received attention in diverse fields of study (e.g., Angrist and Krueger 1999; Aronow and Samii 2016; Angrist and Pischke 2009, pp. 76, 79). Specifically, if we are interested in the average effect of treatment *X* on outcome *Y*, recognizing that this treatment effect likely differs across individuals, but we are also concerned that another variable *Z* confounds the relationship between *X* and *Y*, then, in estimating the multivariate regression

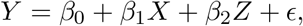

the estimate 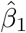 of the treatment effect of *X* is a weighted average of individuals’ treatment effects, with the weight of a particular individual *i* in this average given by

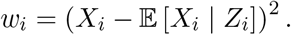

In our case, with reference to Eq. (A.47), *X* = *PGS* and *Z* = *PGS* _*m*_ + *PGS* _*p*_, so that

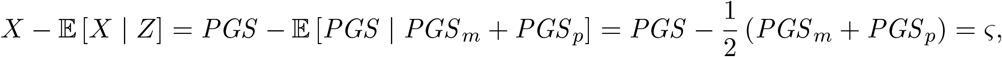

where *ς* is the deviation of the offspring’s PGS from the midparent value, due to the randomness of segregation in the parents. Therefore, estimating the treatment effect of the PGS via least-squares estimation of Eq. (A.47) (or via Eq. A.49) weights individuals according to their values of *ς*^2^, the expected value of which for a given individual is the segregation variance in their family.

Unlike in the more general case where G*×*E effects can differ across loci (Eq. A.54), the estimate of *δ* under the amplification model (Eq. A.57) does have a causal interpretation at the population level. Specifically, if we randomly select an individual from our sample, with probability weighted by their family’s segregation variance, and swap out alleles in that individual so as to increase their PGS by one unit, then 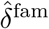 is an unbiased estimate of the average effect on the sampled individual’s phenotype under this procedure. In the terminology of causal inference, 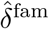 is an estimate of the local average treatment effect (LATE) in an ‘effective sample’ weighted as described above (Aronow and Samii 2016).

In principle, this also implies that we could reweight each family in the least-squares regression, proportional to the inverse of their segregation variance (cf. Aronow and Samii 2016); the estimate produced would then be, in expectation, an average of family-specific slopes *δ*_*f*_, with each family weighted equally.

## Notes

### Competing Interest Statement

The authors have declared no competing interest.

